# Drug-induced adaptation along a resistance continuum in cancer cells

**DOI:** 10.1101/2022.06.21.496830

**Authors:** Gustavo S. França, Maayan Baron, Maayan Pour, Benjamin R. King, Anjali Rao, Selim Misirlioglu, Dalia Barkley, Igor Dolgalev, Kwan Ho-Tang, Gal Avital, Felicia Kuperwaser, Ayushi Patel, Douglas A. Levine, Timothee Lionnet, Itai Yanai

## Abstract

Advancements in rational drug design over the past decades have consistently produced new cancer therapies, but such treatments are inevitably countered through an adaptive process that fosters therapy resistance. Malignant cells achieve drug resistance through intrinsic and acquired mechanisms, rooted in genetic and non-genetic determinants. In particular, recent work has highlighted the role of intrinsic cellular heterogeneity in the emergence of transient drug-tolerant persister cells that survive drug treatment, as well as non-genetically driven cell plasticity toward stable resistance. However, these models do not account for the role of dose and treatment duration as extrinsic forces in eliciting cancer cell adaptation. Here, we show that these two components together drive the resistance of ovarian cancer cells to targeted therapy along a trajectory of cellular adaptation, that we denote the ‘resistance continuum’. We report that gradual dose exposure and prolonged treatment promote a continuous increase in fitness, and show that this process is mediated by evolving transcriptional, epigenetic and genetic changes that promote multiple cell state transitions. The resistance continuum is underpinned by the assembly of gene expression programs and epigenetically reinforced stress response regulation. Using both *in vivo* and *in vitro* models, we found that this process involves widespread reprogramming of cell survival pathways, including interferon response, lineage reprogramming, metabolic rewiring and oxidative stress regulation. Together, the resistance continuum reveals the dynamic nature of cellular adaptation, and carries implications for cancer therapies, as initial exposure to lower doses primes cells over time for increased resistance to higher doses. Beyond cancer, such continuous adaptation exposes a basic aspect of cellular plasticity, which may also be deployed in other biological systems such as development, immune response and host-pathogen interactions.

## INTRODUCTION

The ability of malignant cells to consistently evolve resistance to therapy poses one of the major obstacles to cancer treatment (1). Models to explain the emergence of resistance have traditionally relied upon the selection of genetic mutations that enable cells to escape a drug’s effect (2). More recently the roles of non-genetic mechanisms – in conjunction with genetic changes – have been increasingly recognized in promoting drug resistance (3). Evidence for the effect of epigenetic changes was observed in therapy-induced transcriptional reprogramming of cells primed (4) or equipotent (5,6) to enter in a reversible drug-tolerant state (7). Therapy-induced reprogramming eventually results in heritable adaptive states, imparting stable resistance (8,9). However, by assessing drug resistance as a snapshot in time, these models do not explicitly account for the intensity of the selective pressure (dose) and the duration of the treatment (time) (10), thereby obscuring the dynamics of drug-induced adaptation. Here, we study the effects of dose and treatment duration on the adaptive potential of malignant cells and reveal a landscape of resistant phenotypes that emerge according to the tumor’s treatment history.

## RESULTS

### Dose-escalation facilitates drug adaptation along a resistance continuum

We hypothesized that drug-induced resistance occurs as a gradual process propelled by dose exposure and treatment duration. As an experimental framework, we conceived a long-term dose escalation experiment using a human BRCA2-deficient high grade serous ovarian cancer (HGSOC) cell line (Kuramochi (11)) treated with olaparib – a PARP inhibitor that induces synthetic lethality in BRCA1/2 deficient tumors (12). While olaparib is commonly used clinically, and constitutes the standard of care for treating homologous recombination defective breast and ovarian cancers, the emergence of resistance is pervasive (13). We first seeded the drug-naïve cells at low density (10^6^ cells on 150 mm plates) and treated with 1 μM of olaparib until they reached confluency, thus characterizing adaptation to this dose. The resulting population was then seeded at the same initial density onto a new plate and treated with an escalated dose of 2.5 μM. We repeated this process, each time doubling the drug concentrations, until cells were able to reach confluency at 320 μM, thereby generating a panel of nine adapted cell populations (T1 to T320) over the course of 311 days (Fig. 1a). To compare the dose-dependent responses between these populations – henceforth ‘lines’ – we measured cell viability in each line with the same range of doses used to generate them (1 μM to 320 μM). We found that throughout the dose-escalation treatment, the lines became progressively more resistant, with the half maximal inhibitory concentration (IC50) shifting from ∼2 μM for the control population (C) to ∼60 μM for the cells adapted to 320 μM (T320) (Fig. 1b). This shift depicts an increase in drug tolerance toward higher doses, rather than full resistance, because their survival is higher relative to their parental populations, however even the most adapted lines continue to exhibit sensitivity to the doses to which they had adapted. We also observed that the lines that had adapted to higher doses progressively changed their morphology toward a mesenchymal-like phenotype (Fig. 1a).

**Figure 1.**
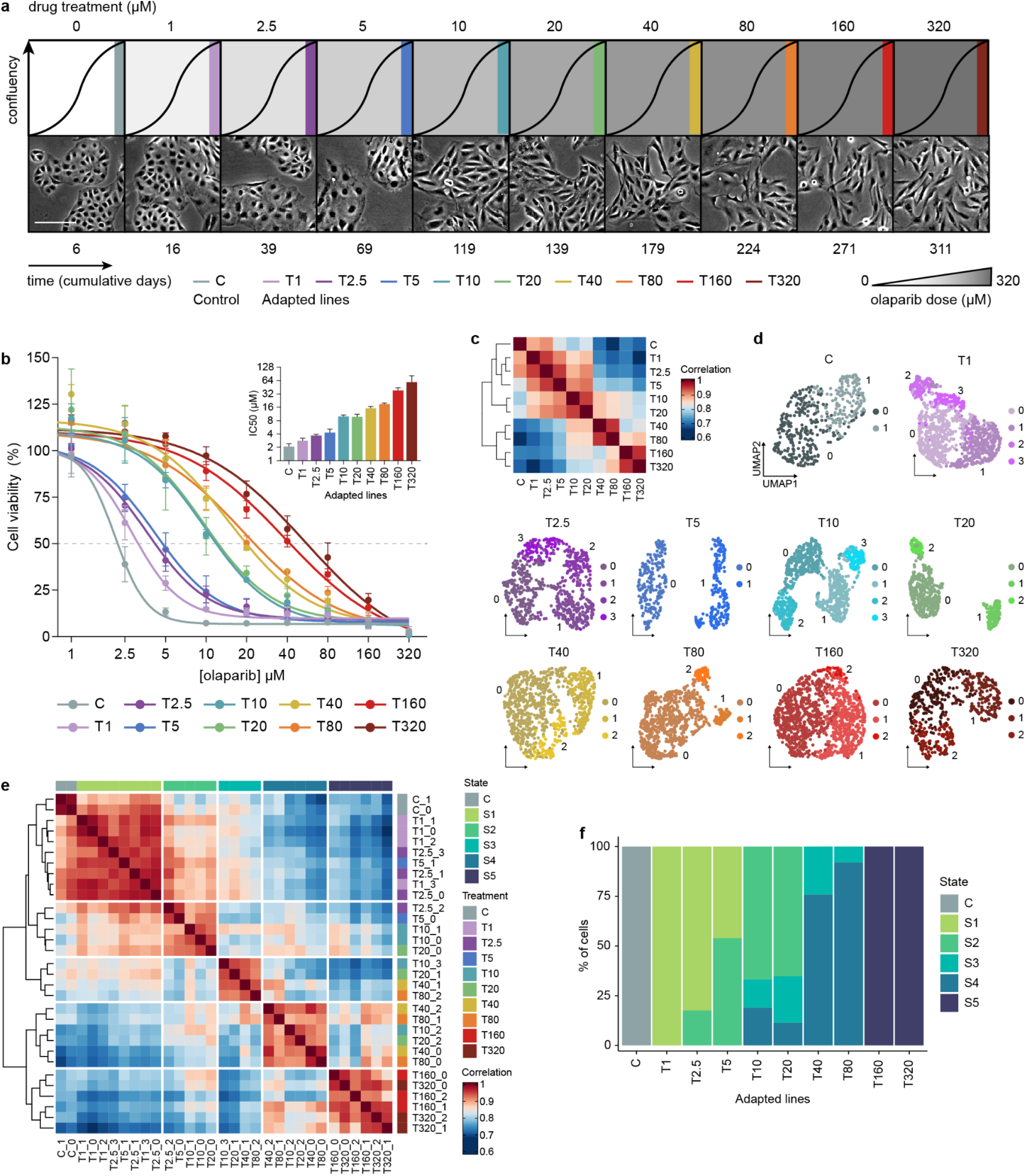
Dose-escalation reveals a progressive fitness increase coupled with multiple cell state transitions. **a.** Schematic of the experimental design for the generation of the drug-adapted lines. The Kuramochi cell line was challenged by increasing drug concentrations (from 1 to 320 μM). Specific doses and duration of treatment are indicated. Cell morphology is shown from representative microscopic images (5x magnification, scale bar = 50 μm). **b.** Cell viability of the adaptive lines showing the olaparib response during 9 days of treatment. The range of doses are the same as those used to generate the lines. All data points are normalized relative to vehicle-treated controls (for each respective line) and represent the average of 3 independent experiments (6 technical replicates per experiment) and their respective standard error bars (s.e.m). **c.** Spearman’s correlations among the averaged transcriptomes of the adapted cell lines. **d.** UMAP representation of scRNA-seq data on the individual lines. Colors and numbers indicate subpopulations determined by Louvain clustering. **e.** Clustering of subpopulations based on their Spearman’s rank correlation coefficients across the adaptive lines. The defined five major transcriptional states are indicated. **f.** Frequency of cells for each population in each of the five states across the adaptive lines. Cells belonging to a particular subpopulation were assigned to their respective states based on their subpopulation clustering shown in Figure 1e.

To exclude the possibility that resistance emerged by the selection of pre-existing resistant clones, we performed two rounds of treatment-recovery experiments on drug-naïve cells and found that the surviving cells reverted to a sensitive state following recovery, suggesting a drug-tolerant persister (DTP) (7) phenotype (Supplementary Fig. 1a). We next asked whether the duration of the treatment could alone elicit the resistant phenotypes, without an increase in drug concentration. To test this, we maintained populations of T5, T20 and T80 cells at the final concentrations to which they had adapted (5, 20 and 80 μM, respectively) for the number of days that they had taken to adapt to their subsequent populations at increased concentrations (T10, T40, T160) (Supplementary Fig. 1b-d). We observed that the prolonged exposure at constant doses did not generate similarly resistant populations for the T5 and T20 extended treatments when compared to their dose-escalated counterparts (T10 and T40), suggesting that dose increase elicits greater resistance. The extended treatments did however lead to more resistant lines relative to their parental populations, either by shifting their IC50s or better survival at the escalated doses (Supplementary Fig. 1b-d). The cells from the T80 extended treatments were similarly resistant to T160 (Supplementary Fig. 1d), thus highlighting the role of treatment duration. Together, these observations indicate that both dose and time contribute to increased drug-resistant phenotypes along the adaptive path.

To address the hypothesis that dose escalation indeed facilitates drug adaptation, we compared the drug resistance phenotypes between cells that were dose-escalated from 1 to 10 μM with drug-naïve cells directly exposed to the same doses for the same amount of time (Supplementary Fig. 1e-h). The emergence of increasingly resistant phenotypes in dose-escalated cells was reproducible across three independent experiments (Supplementary Fig. 1e). Dose-escalated cells were able to proliferate and reach confluency, while drug-naive cells directly exposed to higher doses (5 and 10 μM) did not reach confluency over the same treatment duration, revealing that previous drug exposure to lower doses primes adaptation to drug (Supplementary Fig. 1g,h). Notably, the continuous long-term 10 μM treatment – comparable to a clinically relevant dose (14,15) – led to very few viable cells, which prevented us from testing their fitness with the viability assays. The populations that were dose-escalated up to 2.5 μM were significantly more resistant than their continuous treatment counterparts (Supplementary Fig. 1f).

We also repeated the dose-escalation experiment with two additional *BRCA*-mutant ovarian cancer cell lines (Ovsaho and COV362 (11)) and observed a similar pattern of fitness increase, providing further evidence that dose escalation generates reproducible phenotypic outcomes in cells with diverse genetic backgrounds (Supplementary Fig. 2a-d). Together, our results suggest that gradual drug exposure elicits incremental adaptive responses that unfold along a continuum, as opposed to a transition to a full resistance. We therefore hypothesized that such fitness increase is accompanied by cell state transitions, dictated by cells’ history of dose exposure and treatment duration.

### Emergence of new cellular states recapitulates treatment history

To test whether transitional cellular states accompany the emergence of resistance, we assessed the transcriptional states of the drug-adapted lines using single-cell RNA-seq (scRNA-Seq). Comparing the average transcriptomes across the lines, we observed that treatment history could be recovered – with adapted lines becoming progressively divergent relative to the untreated cells (Fig. 1c, Supplementary Fig. 3a,g for Ovsaho and COV362). To assess transcriptional states at the level of individual cells, we next performed cell clustering on the data from each line. We accounted for confounding effects due to cell cycle by restricting analysis to cells in G1 phase, as assigned by gene expression cell-cycle scoring (16). Cell clustering revealed distinct subpopulations within the lines (Fig. 1d, Supplementary Fig. 3b,c and 3h,i), indicating treatment induced heterogeneity. We wondered whether the emergence of these subpopulations followed a hierarchical pattern, comprising shared subpopulations across the lines. Using a set of differentially expressed genes across the subpopulations and clustering their average profiles, we found that subpopulations across distinct lines clustered together into what we refer to as ‘states’. (Fig. 1e). For example, one subpopulation from the T10 line (T10_1) clusters with a subpopulation from the T5 line (T5_0), while another subpopulation from the T10 line (T10_3) clusters with a subpopulation from the T20 line (T20_1). We note that subpopulations from earlier adaptive steps (e.g., T10_2, T20_2) cluster with subpopulations from later steps (e.g., T40_0, T80_0), highlighting differential intrinsic plasticity within a population toward higher resistance. Examining the change in the frequency of states across the lines reveals the state dynamics as they emerge and become selected over time (Fig. 1f, Supplementary Fig 3d,j). Our results support the notion that drug resistance occurs along a continuum driven by the emergence of multiple new cellular states.

### Assembly of gene module expression accompanies drug-induced adaptation

We next sought to identify the specific expression programs that underpin the identified adaptive states. We implemented a gene module detection approach in which hierarchical clustering is applied to all cells and nested clusters are evaluated according to their differential gene expression (17). The set of the most differentially expressed genes for each particular cluster of cells, relative to the other cells, was used to define gene signatures, and these were then clustered, yielding six gene modules (A-F, see Methods). Comparing cells according to their module expression, we recovered the states defined in the previous analysis (Fig. 2a, bottom).

**Figure 2.**
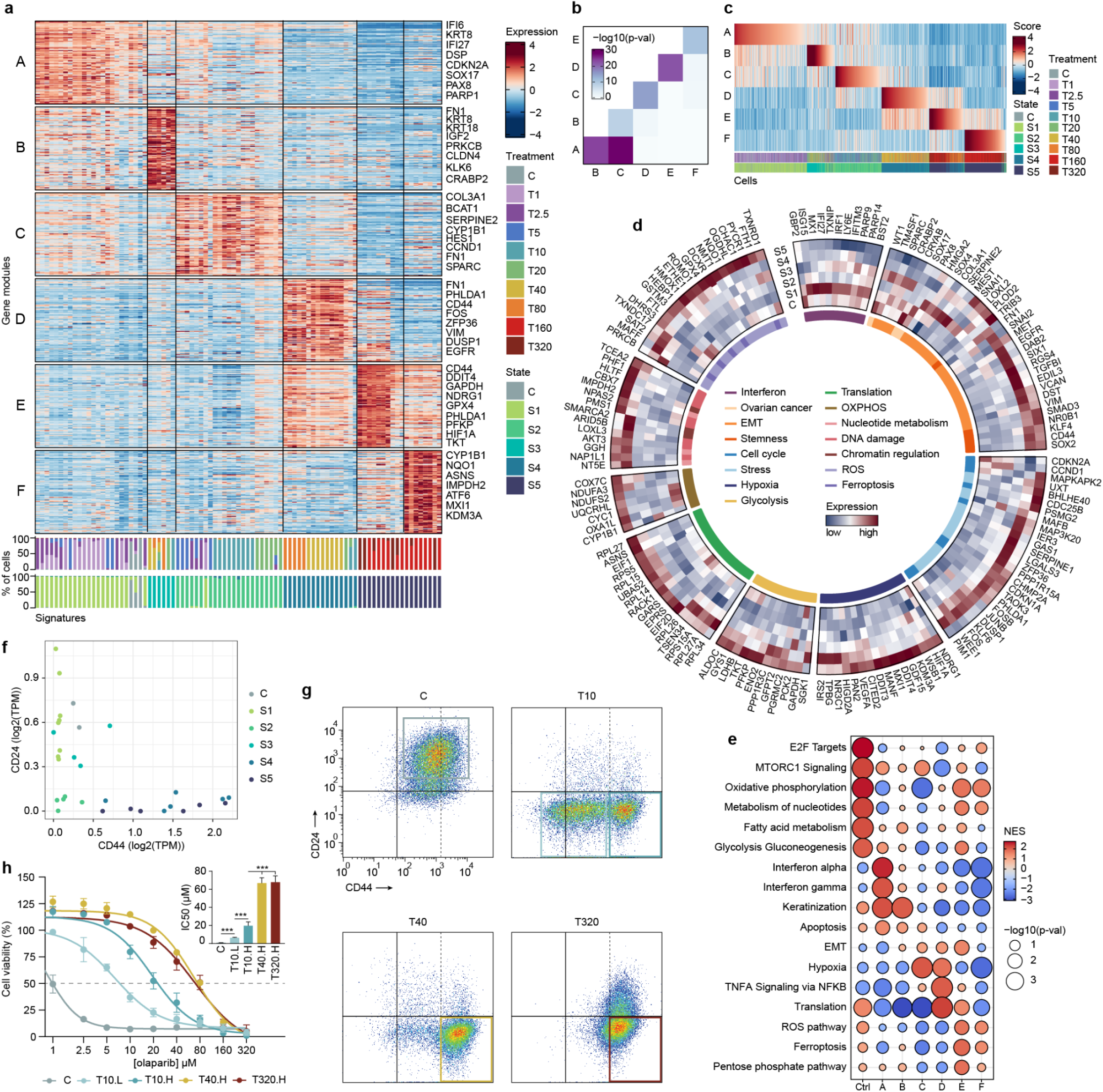
Assembly of gene module expression accompanies drug adaptation. **a.** Gene modules associated with distinct transcriptional states. The heatmap indicates the genes (rows) that are highly expressed in a particular set of gene signatures (columns). Gene signatures are defined as gene sets differentially expressed in a cluster of cells as determined by hierarchical clustering. Each gene module is defined by a set of gene signatures. Barplots (bottom) show for each signature the frequency of cells from the adapted populations and their annotated states. **b.** Overlap of genes across the gene modules showing interdependence in gene module composition. **c.** Cell module scoring. Columns indicate cells from a particular treatment and state sorted by their highest scaled module score. **d.** Gene markers associated with the identified transcriptional states and gene modules, with their annotated functions important for cell survival in drug tolerance. Gene functions were manually curated by literature search in the context of drug tolerance and resistance and annotations from MSigDB hallmark, KEGG and Reactome. Gene expression is represented by the scaled average expression across the states. **e.** Gene set enrichment analysis indicating the relative enrichment of functional annotations for cells that constitute each module. Untreated control cells were considered separately from the module A. Gene sets with *P-*value < 0.01 were selected. Color scale indicates the normalized enriched score (NES) as calculated by GSEA. **f.** Scatter plot showing the average expression of CD24 and CD44 markers in each of the identified subpopulations (d) colored by state. **g.** Flow cytometry analysis recapitulates the transition from CD24^high^CD44^low^ to CD24^low^CD44^high^ in representative populations (C, T10, T40 and T320). The transition across all populations is shown in Supplementary Fig. 4. Colored squares represent the sorted subpopulations used to test for resistance-level differences. Black dashed lines indicate the gating for CD44^high^ populations. **h.** Cell viability assay of sorted subpopulations indicating differential resistance within (T10) and between (T10, T40, T320) treatment conditions. Sorted subpopulations were labeled as “L” (low CD44) and “H” (high CD44). Dots of the cell viability curves represent the average of 2 independent experiments (6 technical replicates per experiment). Bar plot shows differences in IC50 between the sorted subpopulations (\*\*\**P* < 0.001, two-tailed *t*-tests).

The genes present in the six modules comprise a range of molecular functions that may contribute to the survival mechanisms of the drug resistant cells (Fig. 2a, Supplementary Table 1). Cells in state 1 express gene module A, which includes ovarian cancer lineage-defining transcription factors such as *SOX17* and *PAX8* (*18–20*) and the *WT1* prognostic marker (20), as well as epithelial (*KRT8*, *DSP*) and interferon response markers (*IFI6*, *IFI27*), indicating that untreated or lower-dose adapted cells retained their ancestral lineage identity. Cells in state 3 express module B, which has a hybrid expression of epithelial (*KRT8*, *KRT18*) and mesenchymal markers (*FN1*). Cells in state 2 express module C, which includes genes involved in the epithelial-mesenchymal transition (EMT) (*FN1*, *SOX4*, *VIM*) and extracellular matrix (*COL3A1*, *SERPINE2*), but have lower expression of epithelial markers. Cells in state 4 express module D, which also includes a diverse set of EMT genes (*SNAI2*, *VIM*), including mediators of TGF-beta signaling (*SMAD3*, *DAB2*) and markers known to maintain cancer stem cell properties (*CD44*, *KLF4*, *SOX2*) (21–23), suggesting a spectrum of EMT states along the adaptive path as observed in highly metastatic (24) and resistant (25) tumors. The expression of modules E and F by cells in state 5 suggests metabolic rewiring in these highly resistant cells, including the hypoxia master regulator *HIF1A*, glycolytic enzymes (*PFKP*, *GAPDH*) and the pentose phosphate pathway (*TKT*) — which counteracts oxidative stress and supports cancer cell survival in stress conditions (26,27). Consistently, other important genes for oxidative stress regulation controlled by *NFE2L2* (also known as *NRF2*) such as *DCXR*, *NQO1, HMOX1, GPX4, FTL, FTH1* are involved in prevention of DTP cell death (28) through ferroptosis (29) and exert cytoprotective effects from iron toxicity (29,30). Moreover, a number of ribosomal (*RPL14*, *RPS15A*) and mRNA translation genes (*EIF1*, *EIF2D*, *RACK1*) are included in modules D and E, suggesting translation reprogramming — a process controlled by the ATF4 transcription factor through the integrated stress response pathway during adaptation to stress and nutrient deprivation (31) – and is an emerging axis of cell plasticity in tumor progression (32).

The gene module delineation did not enforce mutual exclusivity, and thus a particular gene can be associated with multiple modules. Computing the gene overlap among the modules revealed a sequential pattern of significantly shared genes, which corresponded to cell state emergence (Fig. 2b). For example, module D is expressed in cells from state 4 and overlaps with module C and E, each expressed in cells from states 2 and 5, respectively. Scoring each cell by its module expression further highlighted this pattern: cells scoring highly for a particular module also tended to score highly for the adjacent modules (Fig. 2c). For example, cells highly scoring for module D (mostly state 4 cells), also exhibited high scores for module E (mostly state 5 cells), suggesting that the states build on each other along the adaptive state transitions.

Analysis of the functional annotations captured by our module definitions revealed that a large number of the highly expressed genes along the adaptive path are known to be involved in multiple cell survival programs that confer drug tolerance (33). To put our module genes into the context of these drug tolerance programs, we thus mapped prominent markers across the broad categories of changes in cell identity, interferon response, proliferation and adapting metabolism. This analysis highlighted shared and unique gene expression patterns across drug adaptation (Fig. 2d). Notably, many of these annotated genes were also highly expressed in the most resistant populations of Ovsaho and COV362 cell lines, indicating transcriptional convergence (Supplementary Fig. 3e,k, Supplementary Tables 2 and 3).

To further assess the underlying biological processes overrepresented according to each module, we queried for the enrichment of specific functional gene sets (Fig. 2e). As expected, control cells were enriched for functions related to a proliferative and metabolically active phenotype (E2F targets, oxphos, mTOR signaling) when compared with the adapted cells, which, under drug treatment, slow down proliferation. Module A is enriched for interferon response genes, an endogenous signature induced upon replication stress (34) and DNA-damaging agents (35–37). As previously noted at the gene level analysis, cells related to modules A and B retain epithelial features (keratinization), while a spectrum of EMT enrichment from modules B to E was observed. Consistent with our marker gene analysis, increased resistance was accompanied by extensive metabolic adaptation invoking general stress response programs such as hypoxia, glycolysis, ferroptosis, translation and reactive oxygen species pathways (modules C-F). Again, a similar functional enrichment pattern of highly resistant populations were also observed in our independent experiments using the Ovsaho and COV362 cell lines (Supplementary Fig. 3f,l).

Our analysis suggests that continuous drug adaptation elicits the assembly of interconnected gene modules according to the treatment history of the cells. Furthermore, it led us to hypothesize that subpopulations present within the same line but exhibiting distinct transcriptional states have disparate fitness. To test this, we searched for cell state markers and found that subpopulations from state 1, state 3 and control cells expressed high levels of CD24 and low levels of CD44, while cells from state 2 expressed low levels of both markers. In contrast, cells from states 4 and 5 express low levels of CD24 and high levels of CD44 (Fig. 2f). Using flow cytometry, we validated this shift, revealing a gradual transition from CD24^high^CD44^low^ to CD24^low^CD44^high^ as cells become more adapted to the drug (Supplementary Fig. 4). We then performed cell viability assays on the flow sorted populations from representative lines (C, T10, T40 and T320) in different CD24/CD44 states (Fig. 2g). We found that within the T10 population, CD24^low^CD44^high^ cells (state 4) were indeed more resistant than the CD24^low^CD44^low^ (state 2) ones. Comparing the CD24^low^CD44^high^ subpopulations, we found again that those adapted to higher doses were more resistant (Fig. 2h). This supports the notion that within an individual resistant line, multiple states are present with distinct levels of fitness, while further treatment with increased doses drives higher resistance even among cells in similar states, suggesting ongoing adaptation.

### Persister cell states broadly match the early adaptive states

Given the distinct phenotypic outcomes observed between the adapted (Fig. 1b) and persister (Supplementary Fig. 1a) cell populations, we investigated the differences in the transcriptional programs corresponding to these phenotypes and the effects of acute drug exposure. To test this, we generated two drug-tolerant persister lines treated with 10 μM (P10) and 320 μM (P320) of olaparib, respectively, and compared them with the lines adapted at the same doses (Fig. 3a). This allowed us to assess the differences in gene expression programs induced within days (persisters) versus those emerging on the order of months by dosage escalation. The doses for comparison were chosen based on clinical relevance (14,15) (10 μM; sufficient to decrease viability by 90% in drug-naïve cells, Fig. 1b) and with reference to the highest dose of adaptation in our escalation experiment (320 μM). Interestingly, we observed that cells treated with the highest dose (P320) survived better than those treated with the lower dose (P10) (Supplementary Fig. 5a). Cell cycle analysis showed that the 10 μM treatment caused higher mid-late S phase cell accumulation than the 320 μM (*P* = 0.0006), while the proportion of cells in G1 was higher in the 320 μM condition (*P* = 0.002) (Supplementary Fig. 5b). Cell cycle-related phenotypes are expected as olaparib is known to cause replication stress and DNA damage-associated arrest at G2/M phase (38). This result suggests that extreme doses may trigger a protective response preventing cells from undergoing cell cycle, possibly by off-target effects. Further studies will be necessary to elucidate the mechanisms leading to these apparently paradoxical outcomes.

**Figure 3.**
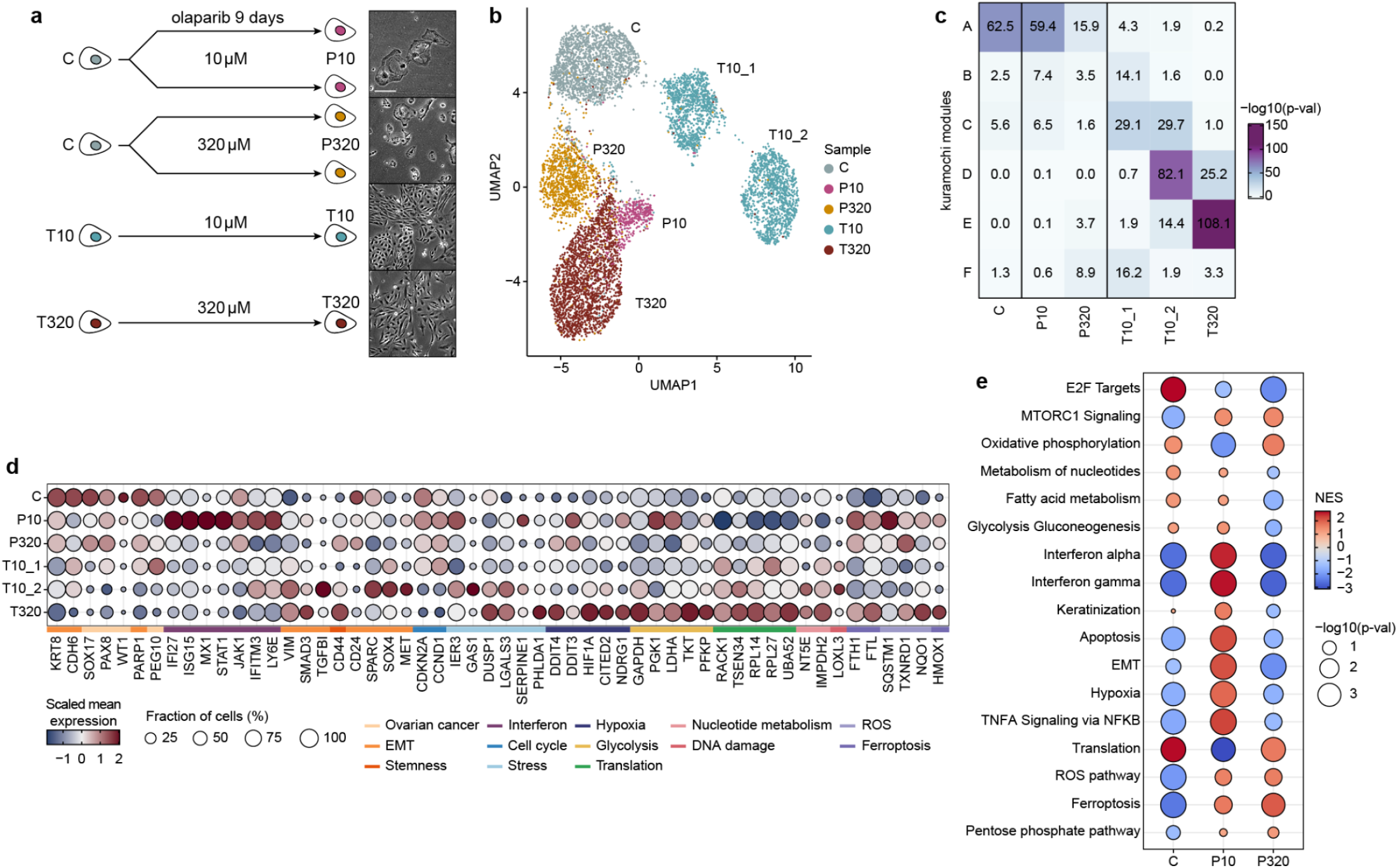
Distinct transcriptional programs for dose-induced persisters and adapted lines. **a.** Drug-tolerant persisters (DTPs) were generated by treating drug-naïve cells with 10 and 320 μM of olaparib for 9 days in two replicates. The adapted T10 and T320 lines were maintained at the same doses. The surviving cells were collected for scRNA-seq. **b.** UMAP representation of the persister and adapted cell populations showing that each condition mainly constitutes a separate cluster. **c.** Gene overlap between the differentially expressed genes across conditions and the previously defined gene modules. Color scale indicates the *P*-values determined from Fisher’s exact tests. **d.** Differentially expressed genes across conditions highlighting shared and distinct marker gene expression among the persister and adapted lines when compared to control cells. Gene functions are labeled according to the annotation depicted in Figure 2d. Color bar represents the scaled mean gene expression for a particular cluster and circle sizes are the fraction of cells expressing the gene. Blue and red indicate low and high expression in a particular condition relative to the others, respectively. Small and big circles indicate low and high percentage of cells expressing the gene, respectively. **e.** Gene set enrichment analysis (GSEA) showing distinct functional enrichment between the persister conditions relative to untreated cells. For the purpose of comparison, the pathways are ordered as in Figure 2e. Color scale indicates the normalized enriched score (NES) as calculated by GSEA.

As expected, although the P10 and P320 lines were treated with the same drug concentrations as the T10 and T320 adapted lines, respectively, we found that they are transcriptionally distinct, indicating a dependence on their treatment history (Fig. 3b). Comparing the overlap of differentially expressed genes in the P10 and P320 with the gene modules delineated for the adapted lines, we observed that the persister states were most related to module A (associated with control and state 1), rather than modules B-F, which were expressed by the highly resistant adapted cell lines (Fig. 3c). Moreover, correlations with the transcriptome profiles that defined the states confirmed these observations, with persisters displaying a higher similarity to the untreated and state 1 cells, while the replicates of adapted lines matched their own state definitions (Supplementary Fig. 5c).

While the extent of transcriptional changes during acute drug response present in the persister lines did not elicit matched adaptive states, resistance markers such as *CD44*, *VIM*, *HMOX1*, *NQO1*, *GPX4*, *DDIT4* and *PFKP,* for example, were highly expressed in persisters relative to untreated cells. Although these genes were expressed at lower levels relative to the adapted lines (Fig. 3d, Supplementary Table 4), their presence suggests an early partial reprogramming, which may prime the subsequent adaptive path. In contrast, the expression of ovarian cancer lineage-dependent factors such as *SOX17* and *PAX8* was retained in persisters (with lower levels relative to C) but lost in the adapted lines (Fig. 3d, Supplementary Table 4). This is consistent with results from other cancer models where differentiation plasticity is coupled with acquired drug resistance (9,39,40), again suggesting that the extent of transcriptional reprogramming is coupled with dose and treatment duration.

Comparing functional gene sets among persisters and untreated cells further revealed processes related to stress response depicted in the adaptive lines (e.g., Interferon, EMT, hypoxia, ROS, ferroptosis) (Fig. 3e). These were particularly enriched in P10, suggesting that treatment with 10 μM elicited a more pronounced stress response compared to 320 μM. This is possibly linked to a higher efficacy of this dose in inducing cell death in the absence of adaptive steps (Supplementary Fig. 5a,b) but also triggering the expression of survival pathways to cope with a higher level of stress (compared to P320), thus highlighting a dose-dependent effect in the generation of persister populations.

### Copy number alterations are associated with the emergence of the adapted cell states

We next asked whether the emergence of our delineated cell states is associated with large-scale copy number alterations (CNAs). We used our scRNA-Seq data to infer CNAs (41) and found that inferred alterations in the adapted lines largely agreed with our defined state subpopulations (Fig. 4a, Supplementary Fig. 6a), indicating a genetic clonal structure in the evolution of the lines. In contrast, clearly distinguishable CNAs were not present among the persister cells (Supplementary Fig. 6a). As independent support for the CNA profiles, we performed whole-exome sequencing (WES) on representative lines (C, T10, T40 and T320) and found concordant patterns (Fig 4a). Interestingly, we found significant associations between the genes present in the modules defined for the adapted lines and their gained or deleted status in the CNA regions (Supplementary Fig. 6b). For example, genes present in deleted regions in T320 (relative to C) significantly overlap with module A genes (lower expression in T320), while genes in the gained regions overlap with genes from modules D and E (higher expression in T320). Notably, some CNAs are conserved across early and late adapted lines (e.g., chromosomes 7 and 9), including a deleted region that spans *CDKN2A* (validated by WES), an important cell cycle regulator recurrently deleted in multiple cancer types and associated with tumor progression (42,43), thus highlighting a potential contribution to PARPi resistance.

**Figure 4.**
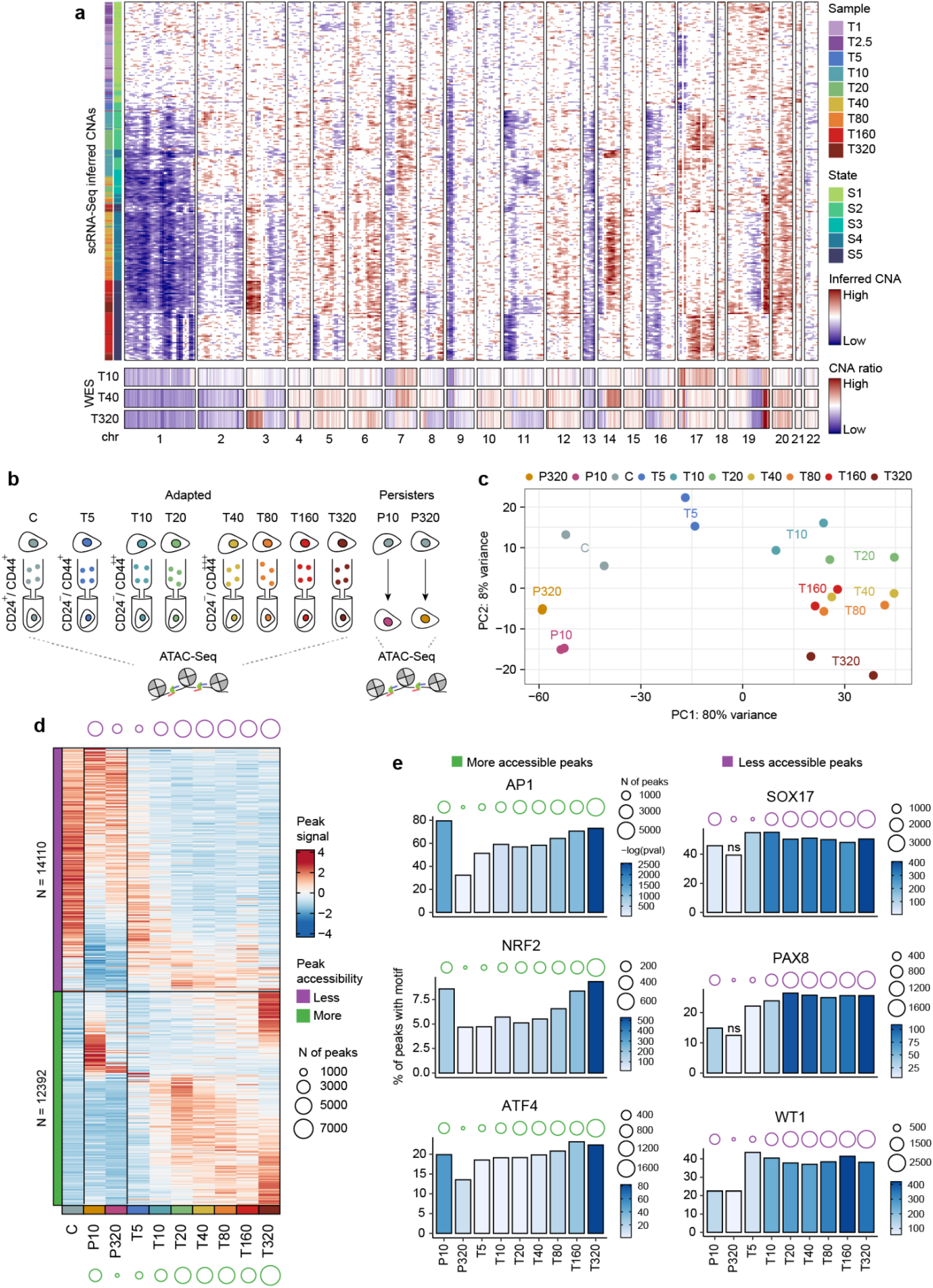
Copy number alterations and epigenetic reprogramming of the adapted and persister lines. **a.** Copy number alteration (CNA) analysis using scRNA-seq inference (top) and whole-exome sequencing (bottom). The top heatmap shows hierarchical clustering of cells depicting the association between cell states and inferred CNAs. Rows indicate inferred CNA profiles for each cell (30% of the cells from each condition were considered for visualization). Whole-exome sequencing of representative lines indicates consistency with inferred CNAs. **b.** Schematic showing the generation of ATAC-Seq data for adapted and persister lines. Cell populations from C and T5-T320 lines were sorted based on their CD24 and CD44 profiles (see Supplementary Fig. 4). Persisters were generated as described in Figure 3a. **c.** Principal component analysis (PCA) on top 2000 most variable peaks across conditions reveals an adaptation trajectory along the PC1 axis. Dots represent data of two replicates for each condition. **d.** Heatmap of the differentially accessible peaks (rows) relative to untreated cells (C). Total number of more and less accessible regions are indicated on the top and bottom parts respectively. Circle sizes represent the total number of differentially accessible peaks for each category per condition. Peaks are ordered based on the method used in Levin *et al*. (48) and shown as scaled values of normalized peak signals. **e.** Transcription factor (TF) motif enrichment analysis. Bar plots represent the percentage of differentially accessible peaks for each condition (relative to C) containing motifs for representative enriched TF families. More accessible peaks in treated conditions are enriched with motifs for stress regulators. Less accessible peaks in treated conditions are enriched with motifs for cell lineage markers. Color bars represent the motif enrichment *P*-values and circle sizes are the number of differentially accessible peaks containing motifs for a particular TF. All significantly enriched motifs are shown in Supplementary Figure 7b.

The association between CNAs and cell states may reflect adaptive changes or they may be a consequence of the PARPi treatment, as replication stress induces CNAs (44,45). Indeed, we observed that the dose escalation experiment across other cell lines (Ovsaho and COV362) led to distinct CNA profiles (Supplementary Fig. 6c), yet convergent transcriptional programs (Supplementary Fig. 3a-l). Moreover, we found that cells belonging to a particular state can exhibit distinct CNA profiles. For example, T160 and T320 cells present in state 5 show distinct inferred CNA profiles (Fig. 4a).

A known genetic mechanism for PARPi resistance is the reversion of the *BRCA1/2* mutations (46). However, examining the single-nucleotide variants (SNVs) from the exome data we did not observe secondary mutations in *BRCA* genes in the resistant lines (T10, T40, T320). These results together suggest that, despite the association between genetic changes and cell states, genetics do not fully explain the emergence of states. We thus next considered the possibility that the emergence of states has an epigenetic component.

### Adaptation-induced chromatin remodeling for stress response and lineage reprogramming

To gain insight into the epigenetic mechanisms that underlie drug adaptation, we measured genome-wide chromatin accessibility by ATAC-seq (47) in the adapted and persister lines. For the adapted lines, we sorted CD24^low^CD44^high^ subpopulations (corresponding to the more resistant cells) from T5 to T320, allowing us to test for continuous epigenetic reprogramming in subpopulations in shared states (Fig. 4b and Supplementary Fig. 4). Examining the chromatin accessibility profiles using principal component analysis (PCA), we observed that PC1 and PC2 together captured the historical order from control cells to T320 (Fig. 4c), indicating an adaptation trajectory of cumulative epigenetic changes and consistent with our transcriptome findings (Fig. 1). Overall, the progression from T5 to T320 was distinguished by an increased number of differentially accessible peaks (relative to C), with shared patterns that suggest heritable epigenetic changes (Fig. 4d). For the persister cells, the chromatin state showed distinct accessibility patterns relative to the adapted lines, again consistent with the transcriptional data (Fig. 4c,d). Of the two persister lines, P10 exhibited a greater number of chromatin changes relative to the P320 (Fig. 4d). We verified that the number of differentially accessible peaks across the lines could not be attributed to peak detection bias (Supplementary Fig. 7a).

We next assessed the enrichment of transcription factor (TF) binding sites among the adapted and persister lines. For a given TF, we queried for an enrichment of its binding sites in the more and less accessible chromatin regions of each sample (Supplementary Fig. 7b). We found that motifs for global stress response regulators AP1 and ATF4 and the oxidative stress sensor NRF2 are enriched in the more accessible regions in the adapted lines. This enrichment progressively increased across the dose-escalation from T5 to T320, in terms of both the higher number of peaks containing such binding sites and their proportion within the differentially accessible peaks (Fig. 4e, left), indicating a reinforcement of those TF stress-regulated programs over time and increased doses. Comparing between the persister lines, we found that P10 showed a higher proportion of peaks containing motifs for those TFs relative to P320 and suggesting that acute exposure at 10 μM, rather than 320 μM, elicits a more exacerbated stress response (Supplementary Fig. 5a,b), in line with our transcriptome observations (Fig. 3d,e). While the P10 and T320 lines exhibited comparable proportions of peaks containing motifs for those stress regulators, the latter accumulated an overall larger number of accessible peaks (Fig. 4e), again highlighting the role of treatment history.

In contrast, peaks containing motifs for lineage-dependent ovarian cancer drivers such as *SOX17*, *PAX8* and the marker *WT1* decreased in accessibility across the adapted lines (in terms of number of peaks), while persisters did not show a similar decrease in accessible regions with such motifs (Fig. 4e, right). Overall, the lower expression (Fig. 3d) and accessibility (Fig. 4e, right) for these TFs indicates a coordinated dedifferentiation program during drug adaptation, a feature that was not observed to a similar extent during acute drug exposure. Of note, we observed significant associations between the differentially accessible peaks and CNAs determined by WES (Supplementary Figure 7d). However, after removing peaks that overlap CNAs, we found that the patterns of TF motif enrichment were not altered, implying widespread epigenetic changes that are functionally independent of CNAs (Supplementary Fig. 7b,c). In summary, these results suggest that drug-induced adaptation is concomitant with extensive chromatin remodeling, whereby stress-responsive regulatory elements are increasingly recruited over dose escalation coupled with a lineage reprogramming-like process.

### *In vivo* analysis of persister and adapted drug-resistant states

We next sought to test for the recapitulation of the persister and adapted cellular states *in vivo*. For this, we designed two sets of experiments using a *BRCA2*-mutant patient derived xenograft (PDX) model of HGSOC (49). In the first set of experiments, we aimed to study whether treatment with different doses elicits a persister-like state. Cohorts of PDX were treated with another PARPi, talazoparib, at four doses (low to high). After 50-57 days of treatment, tumors were collected for scRNA-Seq (Fig. 5a). The corresponding growth curves showed an expected dose-dependent response (Supplementary Fig. 8a). In the second set of experiments, we asked whether prolonged treatment generates PARPi resistance and therefore elicits adaptive-like states. For this, we treated PDX tumors over three serial passages. Vehicles and tumors from the first and third rounds of treatment were collected for scRNA-Seq (Fig. 5b). In a first round, mice were treated with a standard dose (50), which initially caused remission and treatment was interrupted. After an off-treatment period, tumors relapsed and treatment was re-introduced. Tumors did not respond to therapy upon relapse, thus indicating resistance (R1) (Supplementary Fig. 8b). One of the replicates (R1*) was reimplanted for a second round (R2) of treatment and this was followed by a third round (R3), with both under continuous therapy (Fig. 5b). Again, tumors did not respond to therapy in both passages (Supplementary Fig. 8c,d).

**Figure 5.**
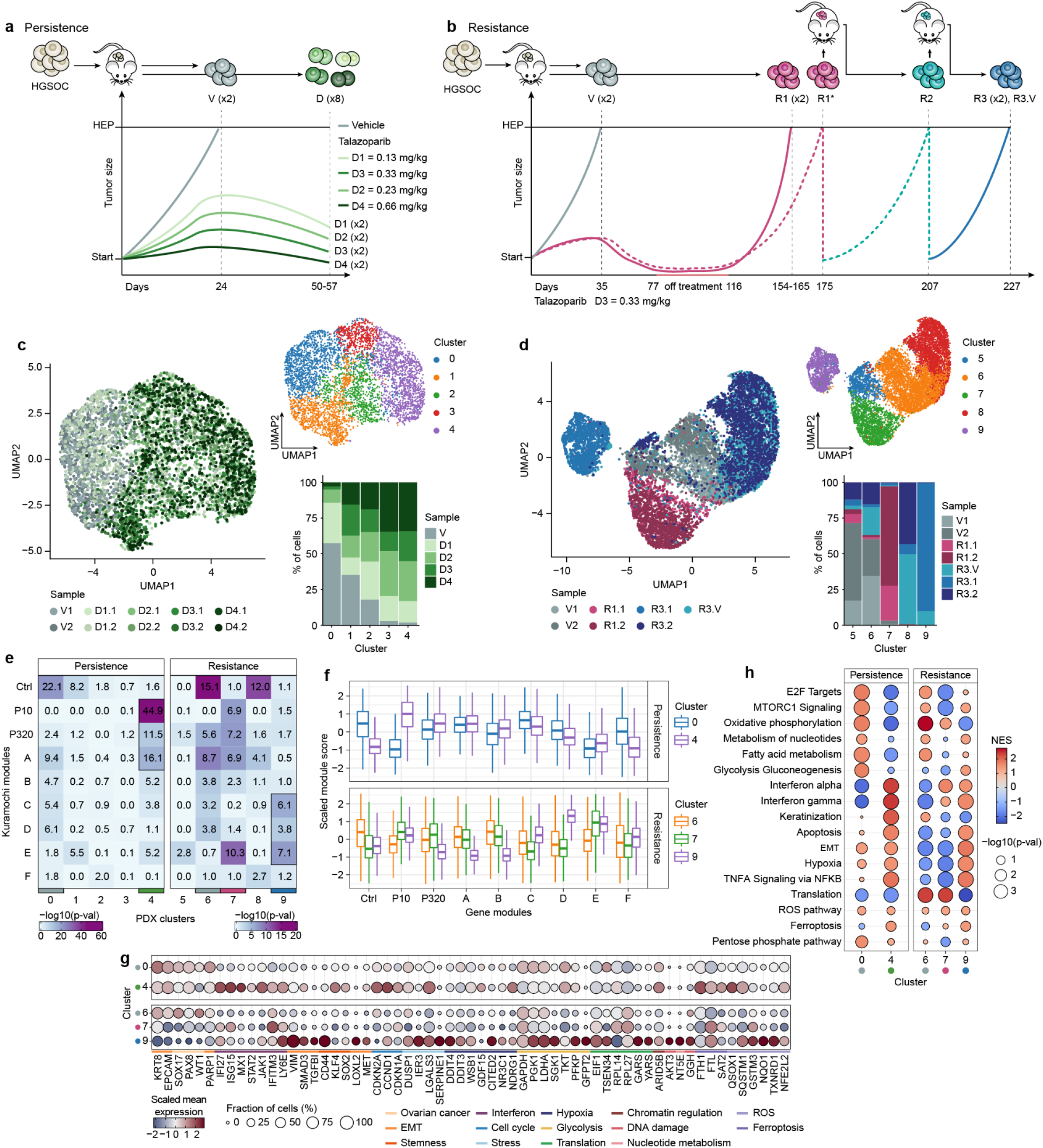
*In vivo* models of dose response and resistance recapitulate the resistance continuum. **a.** Schematic of the experimental design for the PARPi dose-response models of high-grade serous ovarian cancer (HGSOC). Two replicates of each condition (vehicles and talazoparib treated with four doses) were generated. Tumors were collected for scRNA-seq at the humane endpoints (HEP) (vehicles) or after 50-57 days of treatment. The lines represent a schematic of the tumor response. Tumor growth curves are presented in Supplementary Figure 8a. **b.** Schematic of the experimental design for the generation of PARPi resistance models of HGSOC. Three rounds of talazoparib treatment were performed (with a drug holiday indicated in red). Two replicates of vehicle and resistant tumors from rounds 1 and 3 (including a vehicle treated) were collected for scRNA-Seq at the humane endpoint (2000 mm^3^). A replicate from round 1 (R1*) was used to seed the second round of treatment (R2). The lines represent a schematic of the tumor response. Tumor growth curves are presented in the Supplementary Figures 8b-d. **c.** UMAP representation of the Louvain clusters identified across all the conditions and replicates for the persistence experiment. Bar plot indicates the percentage of cells from each condition in each cluster. **d.** Same as described in (c) for the resistance experiment. Bar plot indicates the percentage of cells from each condition in each cluster. **e.** Gene overlap between the differentially expressed genes across clusters from both experiments with the previously defined gene modules. Color scales indicate the *P*-values determined from Fisher’s exact tests. Clusters are highlighted based on their similarity with the untreated controls, persister and adaptive modules. **f.** Boxplots of gene module scores across cells from the persistence and resistance experiments for clusters highlighted in Figure 5e. Scores were determined using the top 100 genes from modules A-F and differentially expressed genes among untreated (C) and persister (P10 and P320) Kuramochi cells. Values show scaled (z-scored) module scores across clusters. **g.** Differentially expressed genes among the clusters with high similarity with the *in vitro* experiments, indicating consistent expression of markers from *in vitro* defined cell states (Figs. 2d and 3d). Gene functions are labeled according to our previous annotations. Color bar represents the scaled mean gene expression for a particular cluster and circle sizes are the fraction of cells expressing the gene. **h.** Gene set enrichment analysis (GSEA) shows consistent functional enrichment for clusters that resemble the persister and the adaptive states. For comparison purposes, the same pathways and order defined from Figure 2e are shown. The enrichment for all clusters are shown in Supplementary Figure 8e.

Analysis of the scRNA-seq data showed that cells from the persistence experiment (dose-response) were largely intermixed (Fig. 5c), while cells from the resistance experiment showed greater heterogeneity with the emergence of condition-specific clusters (Fig. 5d), supporting that treatment duration elicits distinct cell states. Notably, a conspicuous cluster (9) is composed entirely of cells from the tumors of the third round of treatment (Fig. 5d).

To assess whether the *in vivo* experiments recapitulate the state transitions seen in our *in vitro* models, we computed the overlap among the differentially upregulated genes for each cluster with the gene modules from the *in vitro* untreated, persisters and adapted states. Genes from clusters 0 and 6 (composed mostly of cells from vehicle tumors in both PDX experiments) highly overlap with genes from the untreated Kuramochi cells, showing correspondence between the *in vivo* and *in vitro* drug-naïve states (Fig. 5e). We found that cluster 4 genes (from the persistence experiment) overlap more significantly with the modules identified for persisters (P10 and P320) and module A (expressed by the less adapted state 1 Kuramochi cells) (Fig. 5e), indicating a convergent persister-like program. Moreover, genes from cluster 7 (mostly composed of cells from R1 tumors) overlap with persister and module (E) genes, whereas genes that distinguish cluster 9 overlap more significantly with modules C, D and E, indicating convergent adaptive-like programs (Fig. 5e). These findings were supported by scoring the gene modules among cells from these clusters (Fig. 5f).

Examining gene markers from the most resistant lines (states 4 and 5) that are differentially expressed among clusters, we found that they are highly expressed specifically in cluster 9, including mesenchymal (*CD44*, *VIM*, *SMAD3*), hypoxia (*DDIT4*, *CITED2*), glycolysis (*PGK1, PFKP, SGK1*), and oxidative stress markers (*NQO1*, *TXNRD1*) (Fig. 5g and Supplementary Table 5). Again, consistent with our *in vitro* findings, the resistant tumors lost the expression of lineage markers (*SOX17, PAX8, WT1*; clusters 7 and 9), while non-resistant tumors retained their expression (Fig. 5g). We also highlight the recurrent loss of *CDKN2A* expression in the resistant tumors (clusters 7 and 9), possibly a signature of resistance, while the non-resistant tumors exhibited high expression, a signature of growth arrest (51) (Fig. 5g). In agreement with a partial reprogramming observed in the *in vitro* persisters (Fig. 3), tumors from the persistence experiment highly expressed shared markers with the resistant tumors (e.g., *CCND1*, *SQSTM1*, *FTL*, *DUSP1*, *LGALS3*). Increased expression of interferon genes (*IFI27*, *ISG15*, *MX1*, *JAK1*) corroborated an early and acute response to PARPi as observed *in vitro* (Fig. 5g, Supplementary Table 6) and exposes a therapeutic vulnerability with synergistic effects in combination immunotherapy (52,53). In addition, we detected markers that show dose-dependent expression, including genes involved with oxidative stress response (*DCXR*, *QSOX1, SQSTM1*) and its master regulator *NFE2L2* (Supplementary Fig. 8), a signature of persister-cell survival in multiple cancer types (28).

Clusters composed mostly of untreated cells in both experiments (0 and 6) showed higher activity of pathways related to a proliferative and metabolically active state, (e.g., E2F targets pathway, oxphos, mTOR, glycolysis) (Fig. 5h), consistent with our *in vitro* observations. Notably, cluster 4 from the persistence experiment and cluster 9 from the resistant model showed functional convergence, as demonstrated by the higher activity of interferon response, hypoxia, TNFa signaling, apoptosis, EMT and ferroptosis, again reflecting our *in vitro* findings. This corroborates our findings of pathway convergence elicited in both persister and adaptive scenarios. However, the differences in gene expression and module composition between short and long-term treatments underscores the dynamics of transcriptional adaptation over time.

Collectively, the emergence of adaptive states in this clinically relevant model during long-term therapy reveals a pattern that is consistent with the one found during *in vitro* drug adaptation. When tumors were exposed to different doses of PARPi for a shorter period, without the establishment of resistance, we observed a gene expression signature that partially resembles the adaptive states, suggesting that treatment history drives transitions to states of increased fitness along a resistance continuum.

## DISCUSSION

Our study provides a conceptual framework for understanding the pattern and mechanisms by which cancer cells adapt to therapy. We find that increased challenges – in terms of drug concentration and duration of treatment – drive malignant cells to progressive adaptation, a path that we have denoted as the ‘resistance continuum’ (Fig. 6). The resistance continuum unfolds over time in cell populations as a set of state transitions involving distinct functional configurations dictated by both genetic and epigenetic components. This process is akin to the evolution of antibiotic resistance in bacteria, where prolonged and increased drug exposure facilitates the emergence of resistant populations (54). Thus, in contrast with the notion of “fully resistant” phenotypes (3,55–57), we argue that resistance is ongoing and dependent upon the historical events experienced by cell lineages, with dose and treatment duration acting as extrinsic forces in driving the emergence of adaptive states.

**Figure 6.**
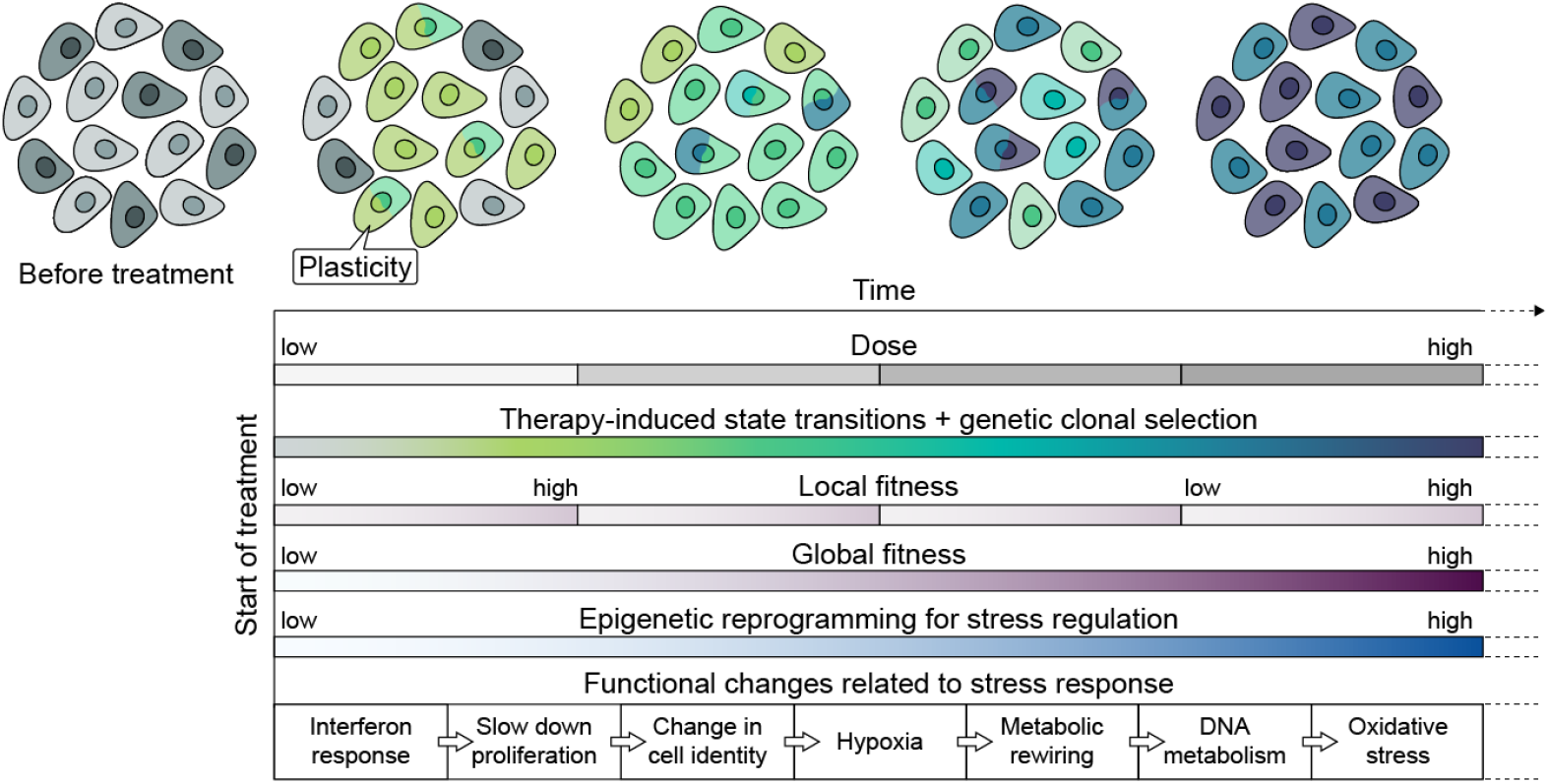
The ‘resistance continuum’ in cancer cell drug-induced adaptation. Continuous treatment coupled with increased doses elicits multiple cell state transitions toward higher fitness. Initial states of drug-naïve cells are represented in gray. Some cells show differential intrinsic plasticity, priming them for state transitions (shades of green and blue) upon therapy. We define fitness as the relative capacity of cells to survive and proliferate under treatment. Cells’ fitness improvement depends on extrinsic forces such as treatment duration (time) and dose exposure. Local fitness gains are achieved from prolonged treatment at constant and tolerable doses. Previous drug exposure and gradual dose increase facilitates adaptation to increasing levels of drug tolerance. Adaptation is the outcome of a complex interplay between the epigenetic reprogramming and genetic changes that affect multiple stress-response programs and resistance mechanisms.

The resistance continuum model is based on the concept that non-lethal drug concentrations induce stress-coping mechanisms that essentially prime cells to assume a more drug-tolerant state. Treatment with higher doses consequently promotes enhanced adaptive states that confer higher fitness with respect to drug resistance. Our model is supported by the observations that progressive transcriptional and epigenetic reprogramming encode multilayered stress mitigation programs upon increasing doses and treatment duration.

This adaptive process has aspects that are both stochastic – due to fluctuations in cell state transitions that define intrinsic plasticity (4,7) – and deterministic. Upon the appearance of a more fit state – either by genetic or epigenetic mechanisms – natural selection acts to fix it in the population, as evidenced by the clonality we observed among the states (Fig. 4a). The process is deterministic in the sense that repeatable phenotypic outcomes (58) – with convergent transcriptional states – are induced by the treatment despite carrying distinct genotypes (Figs. 2, 5 and Supplementary Fig. 3). In contrast to the resistance continuum, drug-tolerant persisters (DTPs) can be viewed as a product of non-adaptive outcomes. For DTPs, a high dose treatment, without previous exposure to milder doses, elicits a “stress endurance mode” (6), but does not necessarily reflect fitness increase in the long term because of their reversible behavior. While some genes or gene programs are partially induced during acute drug response, continuous treatment and gradual dose exposure produces heritable and cumulative changes that appear to facilitate adaptation to stress. In line with this view, a “stochastic tuning” model for cell adaptation to new environments was proposed in yeast (59), whereby the expression levels of individual genes are initially randomly adjusted, and reinforced when resulting in increased cell survival, thus not requiring predetermined genetic mechanisms.

The resistance continuum model provides insight into the adaptive consequences occurring under specific clinical settings, such as intrapatient dose escalation strategies (60,61), short treatment intervals and repeated therapeutic cycles, which may prime cancer cells for adaptation. Moreover, limited drug penetration into solid tumors is a well-known challenge that affects drug diffusion and generates real-life drug gradients (62), due to physical barriers within the tumor architecture (63). The dose escalation dynamics that we studied here may mimic such naturally-created dose gradients, thus exposing cells to tolerable doses and allowing the resistance continuum to unfold over time and space. Our results highlight the need for careful optimization of dose regimen and treatment duration (64,65), coupled with longitudinal monitoring of adaptive outcomes (10,66,67). Determining ‘sweet spot’ combinations of dose and treatment intervals to prevent drug-induced adaptation and persistence may consist of a tangible approach to improve therapy response. Finally, it will be crucial to establish how combination therapies can perturb the resistance continuum’s path for distinct drugs and cancer types by targeting its specific vulnerabilities.

## SUPPLEMENTARY FIGURES

**Supplementary Figure 1.**
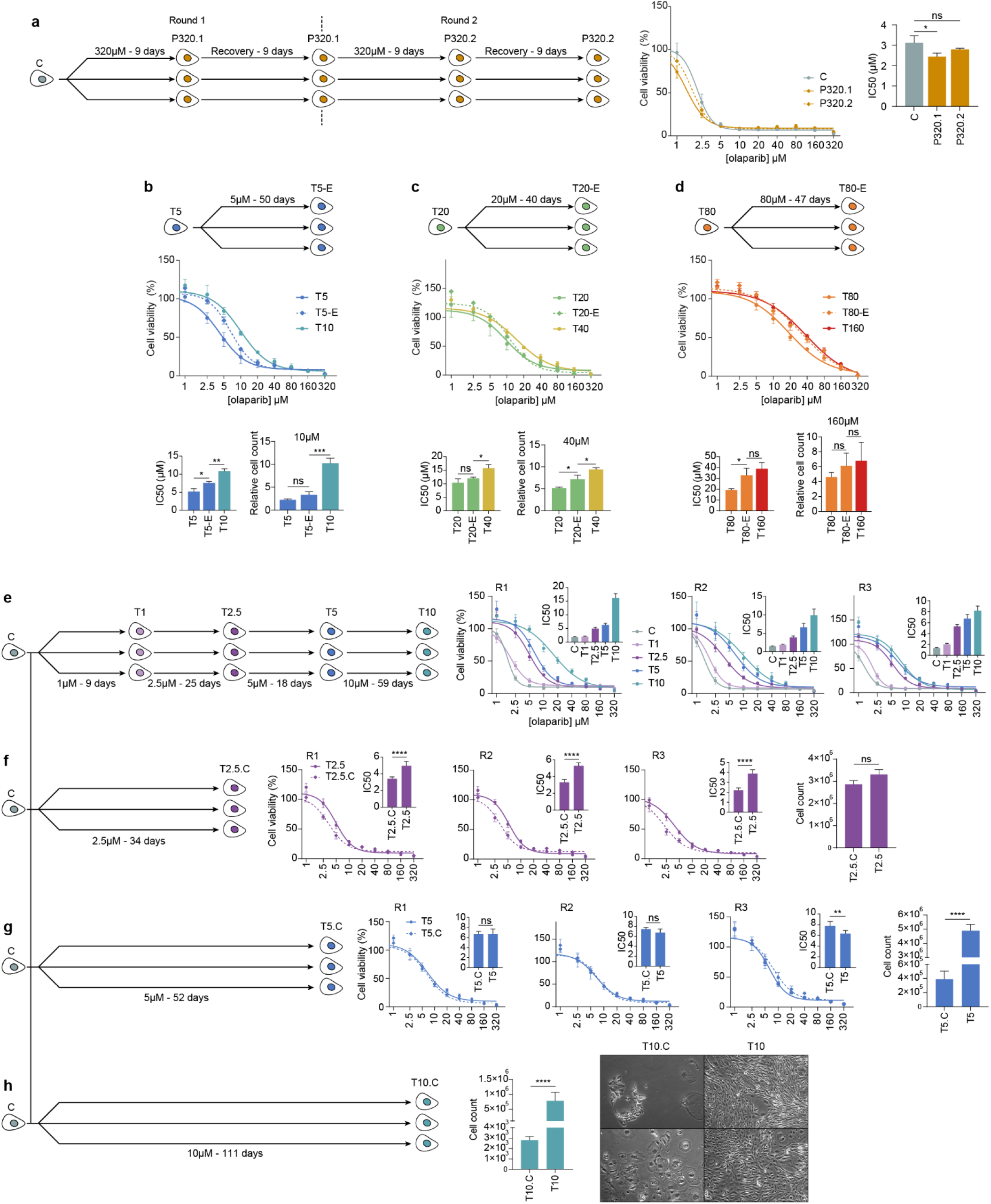
Dose escalation and treatment duration drives the emergence of drug-adapted populations. **a.** Generation of DTPs for Kuramochi cell line by treatment with 320 μM of olaparib for 9 days. The surviving cells were allowed to recover for 9 days without treatment. Two rounds of treatment-recovery experiments were performed in triplicates and the viability of the recovering cells was tested. Cell viability curves are shown as the average of three replicates (n = 6 technical replicates) and their respective standard errors (s.e.m). The bar plot shows no significant IC50 increase for DTPs relative to the parental population (\**P* = 0.03; ns, not significant; two-tailed t-tests). **b‒d.** Extended treatments at constant doses. Representative Kuramochi adapted lines (T5, T20 and T80) were maintained on olaparib treatment at the final concentrations to which they had adapted (5, 20 and 80 μM, respectively) for the number of days that they had taken to adapt to their subsequent populations at increased concentrations (T10, T40, T160, respectively) (see Fig. 1a), generating the extended T5-E, T10-E and T80-E populations. Cell viability was measured by metabolic assays (CellTiter Glo 2.0) and by cell counting after 9 days of treatment at the respective escalated doses (10, 40 and 160 μM). IC50s (left bar plots) and relative cell counts (right bar plots) were compared with their parental and dose-escalated populations (\**P* < 0.05; \*\**P* = 0.005; \*\*\**P* = 0.008; ns, not significant; two-tailed *t*-tests). **e‒h.** Dose escalation facilitates drug-induced adaptation. The dose escalation experiment (Fig. 1a) was repeated in parallel for three replicates until Kuramochi cells reached confluency at 10 μM (e). Plots indicate cell viability curves and shifts in their IC50 (bar plots). Parallel experiments were performed without dose escalation, maintaining drug-naïve cells under continuous treatment at 2.5 (f), 5 (g) and 10 μM (h) for the same duration as the escalation experiments. Cell viability assays and cell counts after the treatment periods are shown and compared with the dose-escalated populations (\*\**P* < 0.01; \*\*\**P* < 0.001; \*\*\*\**P* < 0.0001; ns, not significant; two-tailed *t*-tests). Microscopic photos depict T10 (dose-escalated) cells reaching confluency while T10.C (continuous) did not.

**Supplementary Figure 2.**
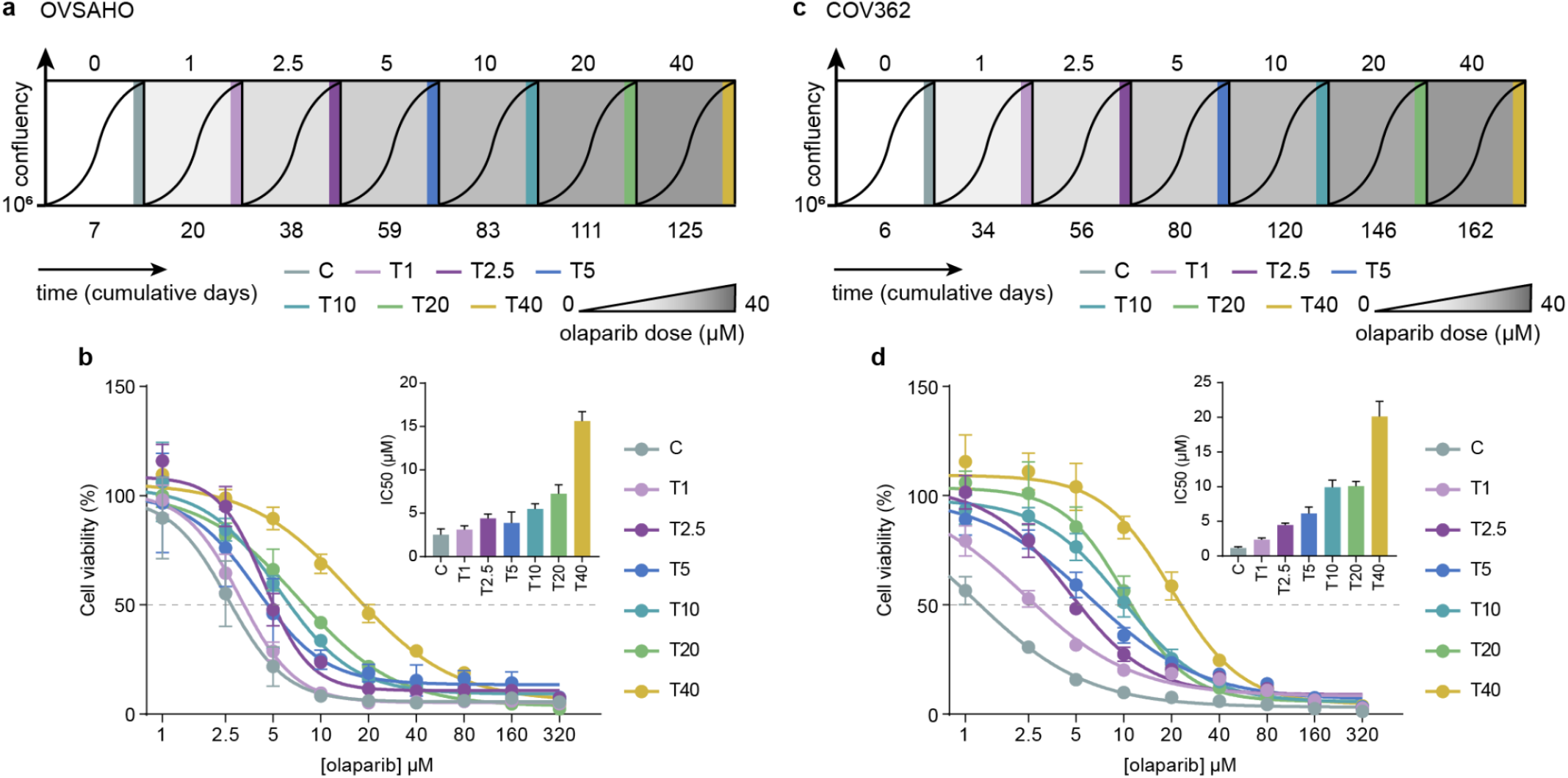
Generation of drug-adapted populations for two additional BRCA mutant ovarian cancer cell lines. Drug-adapted lines were generated using the same dose escalation approach from Figure 1a until cells were able to reach confluency at 40 μM of olaparib. The experiment was performed for Ovsaho **(a)** and COV362 **(c)** and their respective shifts in IC50 are shown by cell viability experiments (**b**, **d**). Data are the means and s.e.m of 6 technical replicates.

**Supplementary Figure 3.**
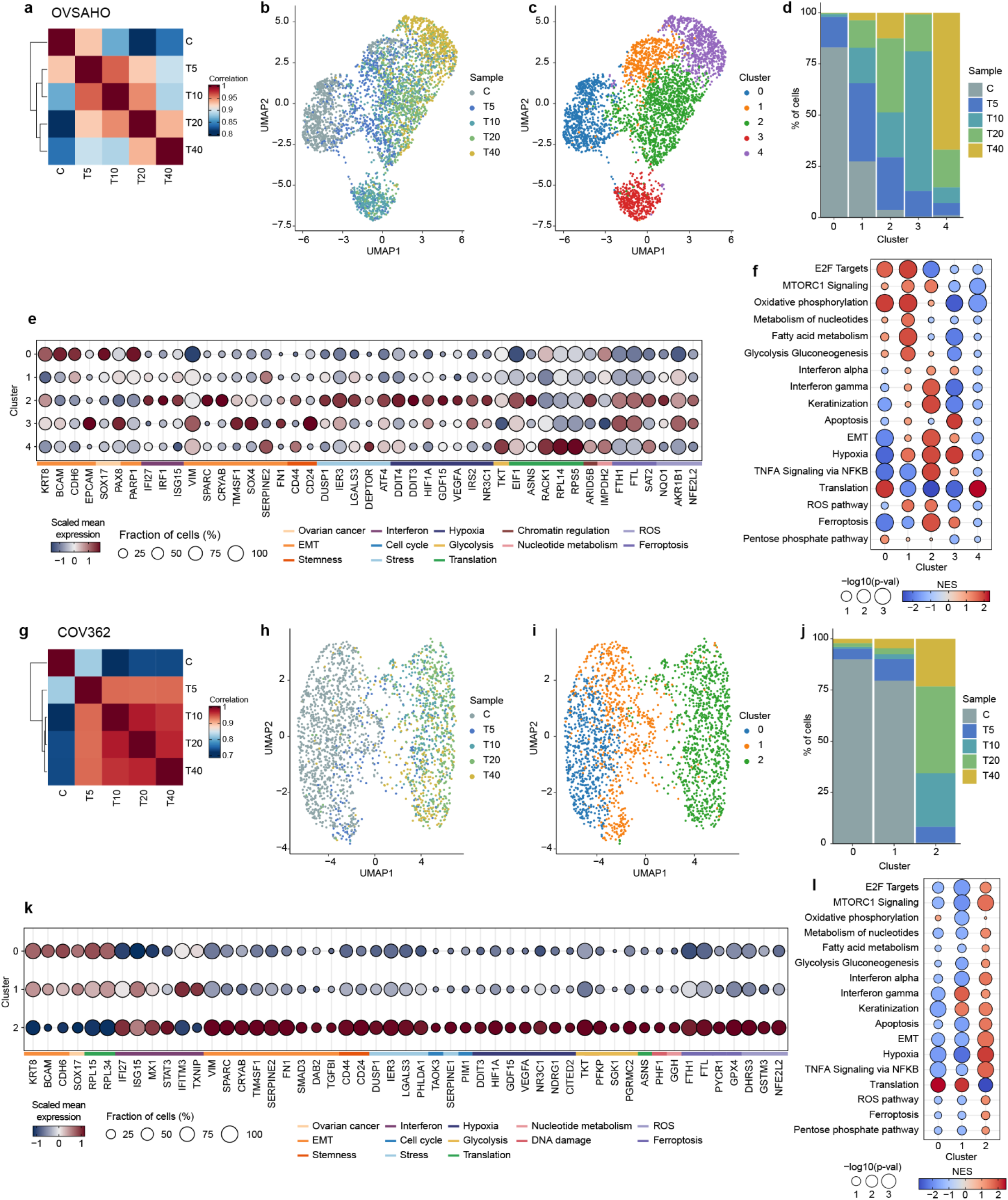
Gene expression in the Ovsaho and COV362 adapted lines. **a.** Spearman’s correlations among the averaged transcriptomes of the Ovsaho adapted lines. Untreated control and T5 to T40 were collected for scRNA-Seq. **b-c.** UMAP representation of Ovsaho scRNA-seq data on the individual adaptive lines colored by treatment (b) and by cluster as determined by Louvain clustering (c). **d.** Bar plot representing the percentage of cells from each condition in each cluster. **e.** Differentially expressed genes indicating consistent patterns of marker expression with previously defined cell states and functional annotation (Fig. 2d). Color bar indicates the scaled mean gene expression for a particular cluster and circle sizes are the fraction of cells expressing the gene. **f.** Gene set enrichment analysis (GSEA) shows consistent functional enrichment for clusters that resemble less (cluster 0) and more adapted (cluster 2, 3, 4) states. For the purpose of comparison, the pathways are ordered as in Figure 2e. **g-l.** Same analyses as shown in a-f for COV362.

**Supplementary Figure 4.**
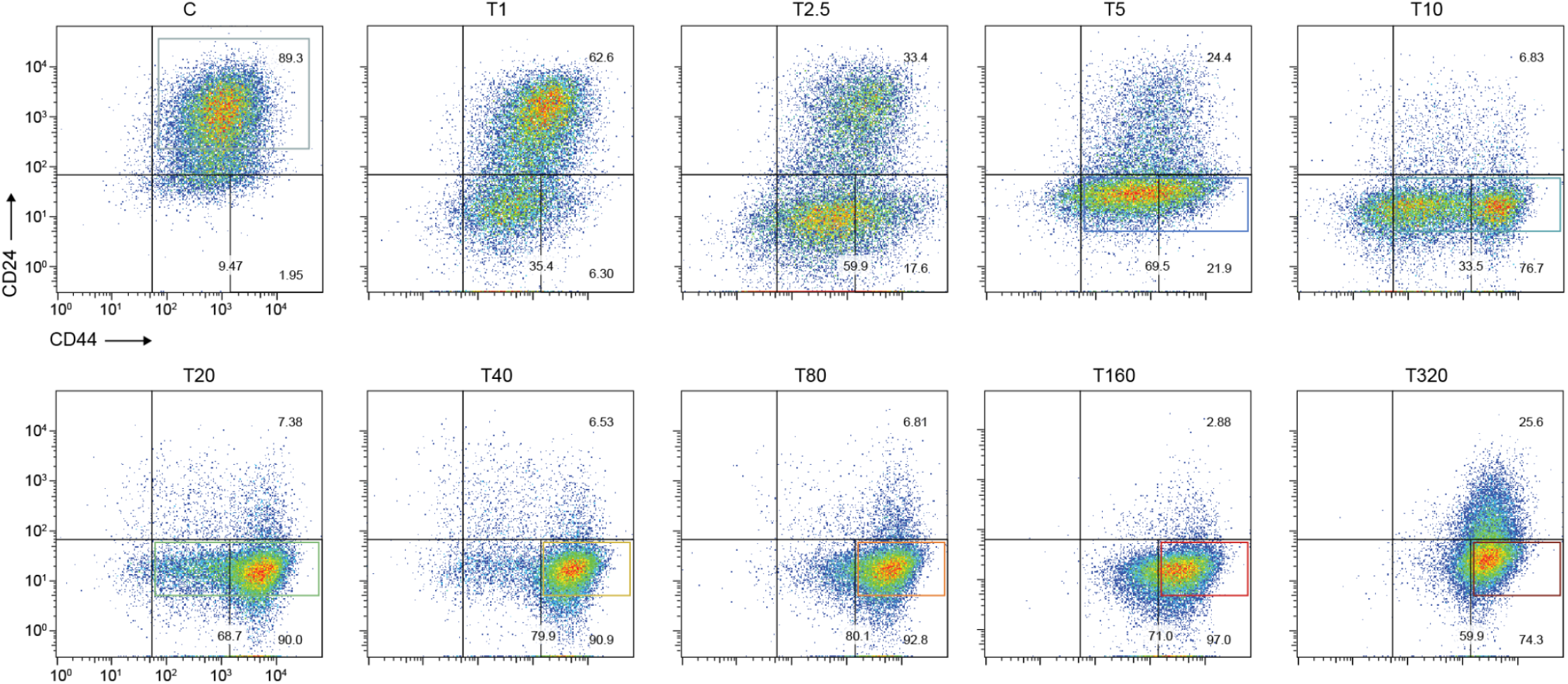
Representative plots of the flow cytometry analysis of CD24 and CD44 staining across the Kuramochi adapted populations. Bottom left quadrants represent double negative staining for both markers. A transition from CD24 ^high^CD44^low^ (top right) to CD24^low^CD44^high^ (bottom right quadrants) is observed. Numbers indicate the percentage of cells in each quadrant. Colored squares indicate the subpopulations sorted for ATAC-Seq (Fig. 4).

**Supplementary Figure 5.**
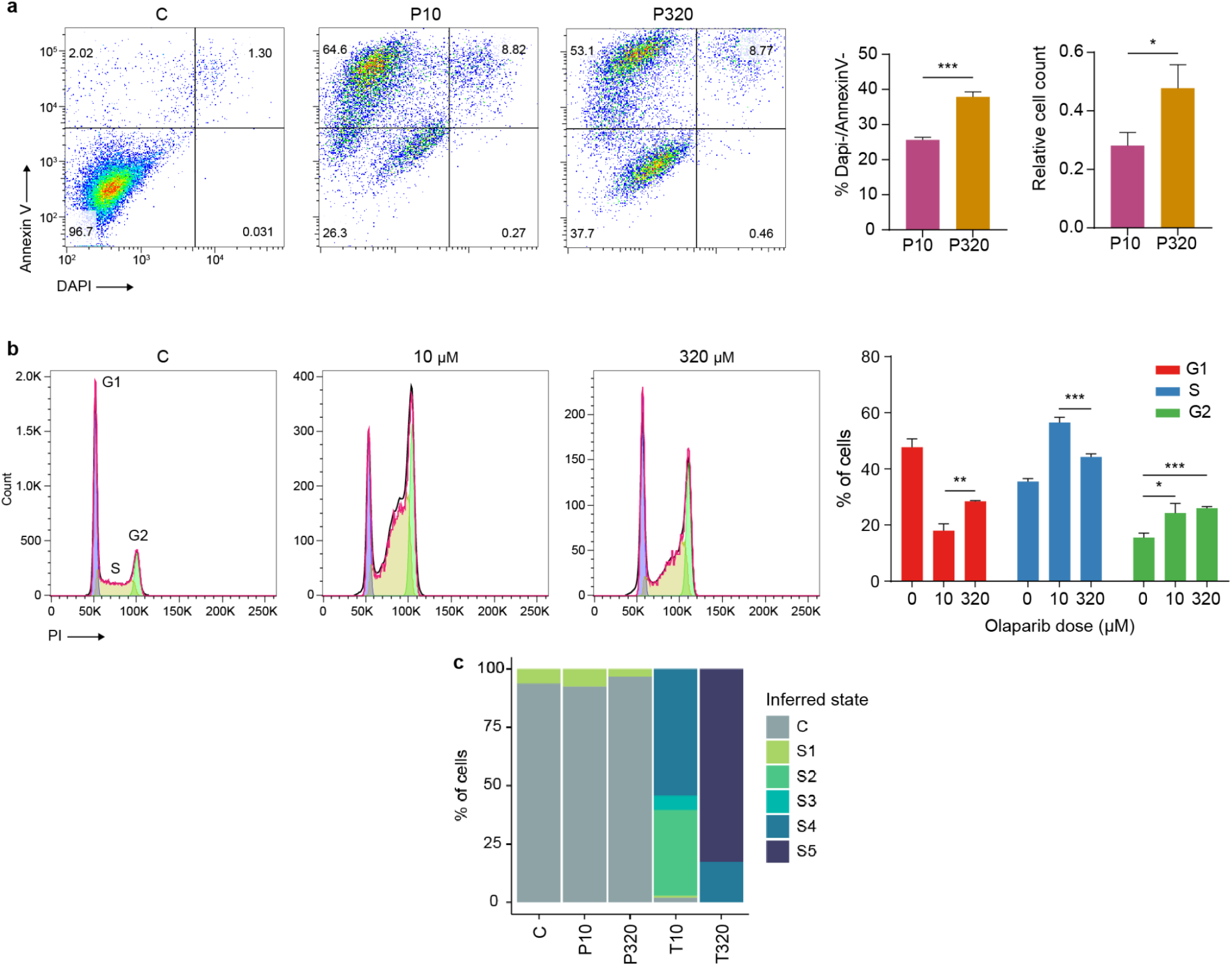
Dose-induced drug tolerant persisters. **a.** Apoptosis assay on cells treated with 10 and 320 uM of olaparib for 9 days. Viable cells were considered double negatively stained for Annexin V and DAP (bottom left quadrant). Percentages of DAPI and Annexin V negatively stained cells and the total number (relative to the initial 1.2 × 10^6^ seeded cells) of viable cells (counted by trypan blue exclusion) are shown as bar plots (\**P* = 0.01, \*\*\**P* = 0.001, two-tailed *t*-tests; error bars are s.e.m of three replicates). **b.** Cell cycle analysis after 2 days of treatment with 10 and 320 uM of olaparib. Histograms represent the cell count in each cell cycle phase. Bar plots represent the fraction of cells in G1, S and G2/M phases (\**P* = 0.01, \*\**P* = 0.001, \*\*\**P* = 0.0001, two-tailed *t*-tests). **c.** Frequency of cells presenting the highest transcriptome correlation with the average profile defined for the adaptive states (Fig. 2).

**Supplementary Figure 6.**
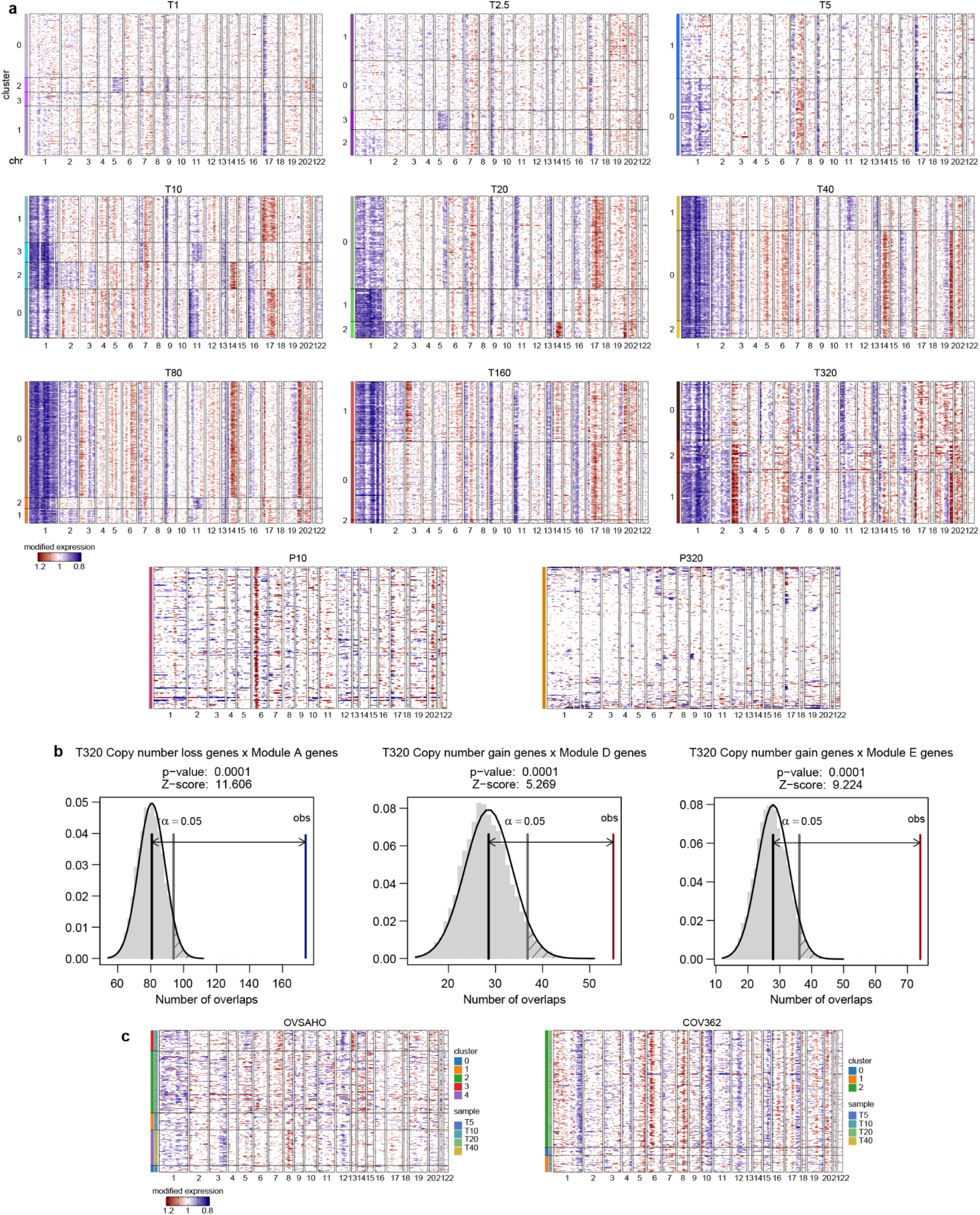
Single-cell CNA inference analysis for the adapted and persister lines. **a.** Heatmaps depicting scRNA-Seq inferred copy number alterations (CNAs) for Kuramochi adapted and persister lines relative to untreated control (C). Gene expression signal is scaled along the chromosomes as calculated by inferCNV. Red represents genomic regions with inferred copy number gain and blue represents regions with inferred copy number loss. Colors and labels on the left represent the subpopulations identified by clustering cells based on their transcriptomes (Fig. 1d) showing correspondence between inferred CNAs and transcriptional states. **b.** Enrichment of genes within CNA regions identified by whole-exome sequencing overlapping with the genes present in the transcriptional modules detected for the adaptive lines (Fig. 2) in respect to random distributions (*P*-values determined by 10,000 permutations, see Methods). Blue and red lines indicate the observed overlap of copy number loss and gain, respectively, and the gray lines indicate the significance threshold in an expected random distribution. **c.** Inferred CNAs defined for the adapted Ovsaho and COV362 lines relative to their untreated controls. Color annotations represent sample and transcriptional clusters identified in Figure S3.

**Supplementary Figure 7.**
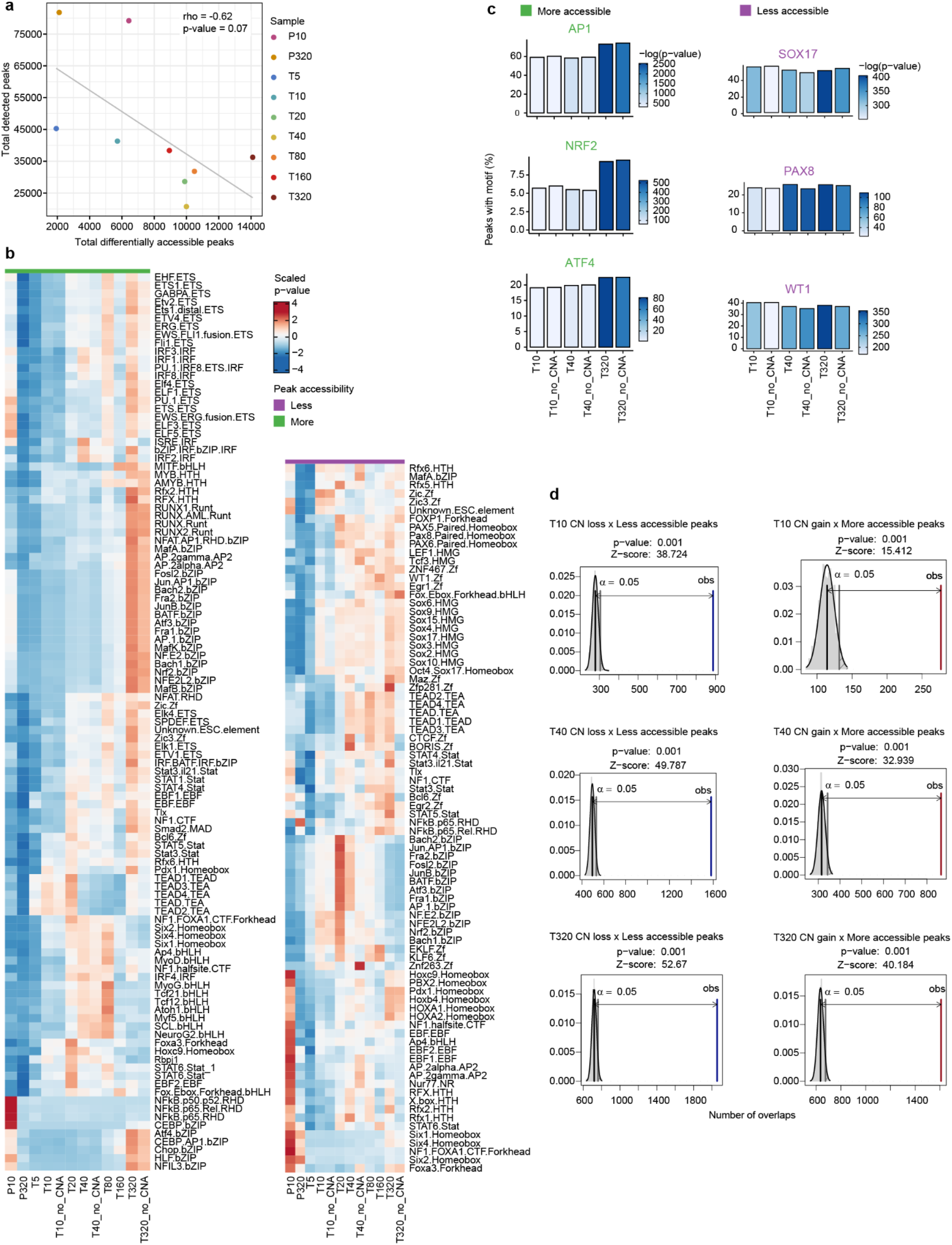
Transcription factor enrichment analysis for the differentially accessible ATAC-Seq peaks and associations with CNAs. **a.** Correlation between the number of differentially accessible peaks and total number of detected peaks per sample. The lack of a positive correlation indicates that the number of differentially accessible peaks are not explained by peak detection bias (Spearman’s rho = −0.62, *P* = 0.07). **b.** Heatmaps showing the hierarchical clustering (rows) of the enriched transcription factor binding sites in the differentially accessible peaks. Green represents TFs with binding sites in peaks with higher accessibility in the adapted lines relative to the untreated condition, and purple for peaks with lower accessibility in the adapted lines. Color bar represents the scaled p-values for transcription factor binding site enrichment (only transcription factors with *P*-value < 1e-5 were included). The analysis controlled by peaks not overlapping CNAs (“no_CNA”) reveals the same patterns when compared to all peaks. **c.** Transcription factor (TF) motif enrichment analysis controlling for peaks not overlapping with CNAs, showing the same frequency of peaks containing motifs for specific TFs involved with regulation of general stress response (*AP1*, *NRF2*, *ATF4*) and ovarian cancer TFs (*SOX17*, *PAX8*, *WT1*). Bar plots represent the percentage of differentially accessible peaks for each condition (relative to C). Color bars represent the motif enrichment *P*-values. **d.** Enrichment of differentially accessible ATAC-Seq peaks overlapping CNA regions (relative to untreated control) identified by whole-exome sequencing for representative adaptive lines (T10, T40 and T320) (*P*-values determined by 1,000 permutations, see Methods). Blue and red lines indicate the observed overlap of copy number loss and gain, respectively, and the gray lines indicate the significance threshold in an expected random distribution. The proportion of peaks not overlapping CNAs ranged from 70% for less and 80-90% for more accessible peaks, highlighting widespread global chromatin changes.

**Supplementary Figure 8.**
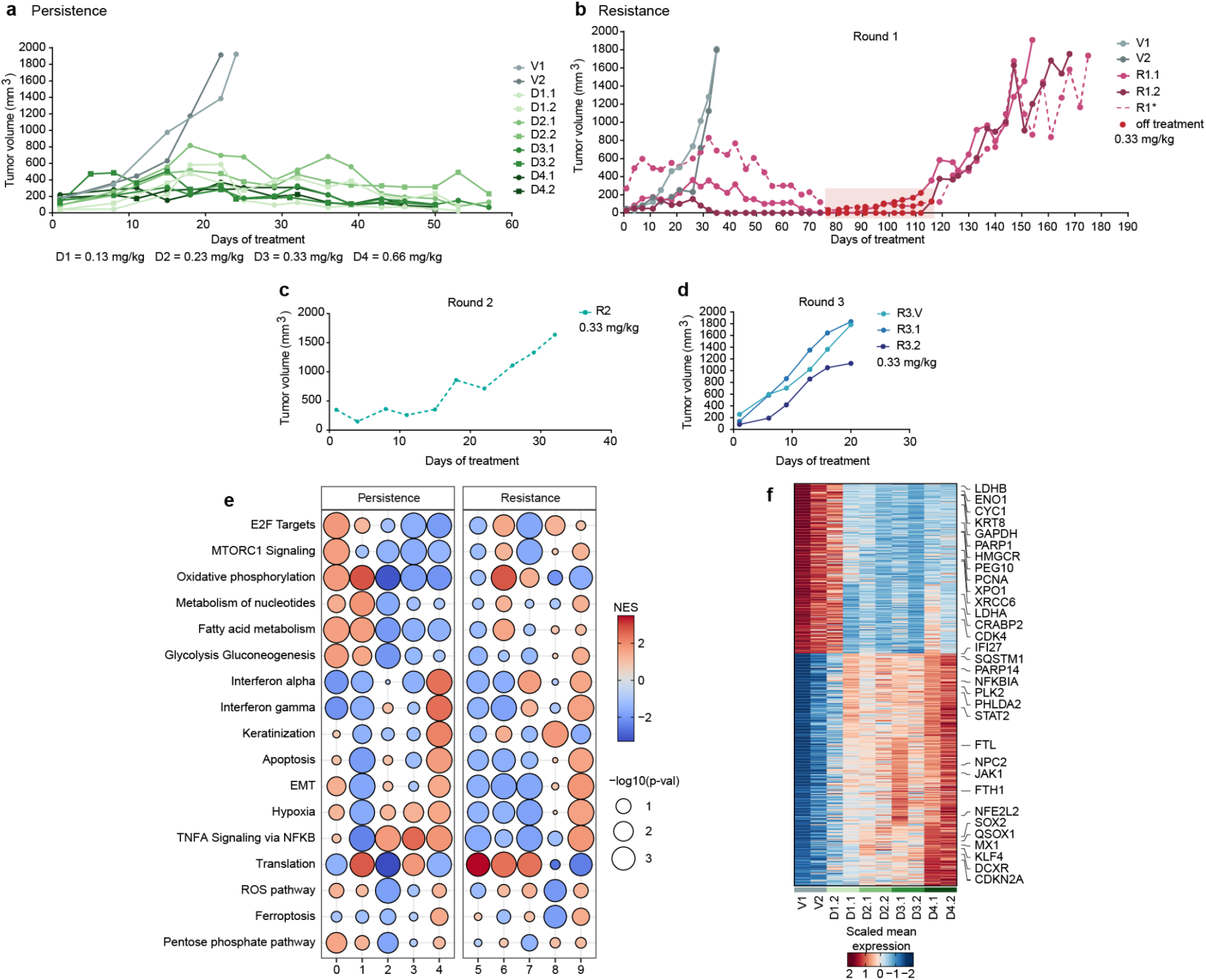
Tumor growth curves and transcriptome analysis of ovarian cancer patient-derived xenografts. **a.** Tumor growth curves for PDXs treated with four doses of talazoparib (shades of green) and vehicle controls (gray). Each condition was collected in duplicates for scRNA-Seq. Two replicates of each condition were generated (indicated by “.1” and “.2”). **b-d.** Tumor growth curves for PDXs in first (b), second (c) and third (d) rounds of talazoparib treatment for the emergence of resistant tumors. A drug holiday in the first round is indicated in red. Two replicates for each condition were collected for scRNA-Seq: vehicles (gray), round 1 (solid pink) and round 3 (blue, with an additional vehicle treated (R3.V)). A replicate tumor from round 1 (dashed pink) was seeded in a new cohort to generate the round 2 treatment. **e.** Gene set enrichment analysis (GSEA) for all the clusters identified by Louvain clustering in both experiments showing convergent signatures with persistence (dose-response) and adaptive states (resistance). For comparison purposes, the same pathways and order defined from Figure 2e are shown. **f.** Heatmap of the scaled gene average expression across samples of the dose-response experiment, highlighting differentially expressed genes that show a dose-dependent pattern.

## METHODS

### Cell culture

Kuramochi cells (*BRCA2*-mutant (11)) were cultured in RPMI-1640 (ThermoFisher) with 10% Fetal Bovine Serum (FBS), 1% MEM Non-Essential Amino Acids, 1% L-Glutamine 200 mM, 1% Antibiotic-Antimycotic solution 100X and 0.25 U/mL insulin solution (Sigma-Aldrich). Ovsaho cells (*BRCA2* homozygous deleted (11)) were cultured in RPMI-1640 (ThermoFisher) with 10% FBS, 1% L-Glutamine 200 mM and 1% Penicillin and Streptomycin (10,000 U/mL, ThermoFisher). COV362 cells (*BRCA1*-mutant (11)) were cultured in DMEM (ThermoFisher) supplemented with 10% FBS, 1% L-Glutamine and 1% Penicillin and Streptomycin. Cells were maintained at 37°C with 5% CO_2_. All cell lines were kindly provided from the laboratory of Benjamin Neel (NYU Langone Health). The cell line identities were confirmed by genotyping using short tandem repeat fingerprinting and were tested negative for mycoplasma with Mycoplasma PCR Detection Kit (ABM).

### Dose escalation experiments

To generate drug-induced resistant populations, 1×10^6^ drug-naïve cells were initially seeded in 150 mm plates. Twenty four hours after seeding, 1 μM of olaparib (Selleckchem, S1060) was added and cells were maintained under treatment until reaching confluency (> 70%), thus characterizing resistance at this dose. Cells were harvested (0.25% Trypsin/EDTA for 5 min at 37°C) and a fraction of the surviving population (1×10^6^ cells) was seeded again and treated with 2.5 μM until cells reached confluency. This process was repeated sequentially by doubling the drug concentrations until the cell populations were able to achieve confluency at 320 μM of olaparib (see schematics from Fig. 1a and Supplementary Fig. 2). The initial dose was determined by cell viability assays (see below), indicating a low starting dose (< IC30). At each step, aliquots of the adapted populations were frozen (10% DMSO, 50% FBS, 40% media) for further experiments. Using this approach, we generated 9 drug-adapted lines for Kuramochi (T1-T320) and 6 adapted lines for Ovsaho and COV362 (T1-T40) plus the untreated controls (C).

To compare the resistant phenotypes generated by dose escalation versus constant dose treatment, we repeated the dose escalation in triplicates for Kuramochi up to 10 μM, while initially drug-naïve cells were maintained under treatment with the same doses (1, 2.5, 5 and 10 μM) for the same amount of time in each step of the dose escalation experiment (see Supplementary Fig. 1e-h for schematics). Media with fresh drug was replenished every 3 days. Olaparib was added 24 hours after plating, in the final concentrations from a stock solution of 50 mM dissolved in DMSO and maintained at −80°C. Cell number was determined by trypan blue exclusion (Sigma-Aldrich) and counted on a Countess II cell counter (Life Technologies).

### Cell viability assays

Viability experiments to test the resistance phenotypes (here defined as fitness) were performed using the CellTiter-Glo 2.0 Luminescent Cell Viability Assay (Promega) according to the manufacturer’s protocol. Before performing the viability assays, frozen aliquots of the resistant cells were thawed and cells were recovered without drug treatment for 7-10 days. Cells were plated in 96-well plates at seeding density of 700 to 800 cells per well. After 24 hours, cells were treated with olaparib with the same concentrations used in the dose escalation experiment (1 to 320 μM) for 9 days. DMSO without drug was used as a negative control to normalize the cell viability of drug treated wells. Luminescence was determined using the BioTek Cytation 5 Cell Imaging Multi-Mode Reader (Agilent). Cell viability analysis was performed on GraphPad Prism 9 and IC50 values were determined using a nonlinear regression curve fit. Pairwise significant differences between IC50s were assessed by Student’s two-tailed *t* tests.

### Cell cycle analysis

Cells were treated with 10 μM or 320 μM of olaparib 24 h after plating and cell cycle analysis was performed after 48 h of treatment using propidium iodide staining. Cells were fixed in cold 70% ethanol overnight at 4°C, washed twice in PBS and resuspended in 1x PBS with 0.2 mg/ml RNase A (Sigma-Aldrich) and 25 mg/ml propidium iodide (Biolegend). After 30 min incubation at room temperature in the dark, cells were analyzed on a LSR II Flow Cytometer (BD Biosciences). Quantification of cell cycle phases was performed on FlowJo software (FLOWJO).

### Drug-tolerant persister generation

Persister cells (DTPs) were generated similar to Sharma *et al*. (7) and Hangauer *et al*. (68) by seeding 1×10^6^ Kuramochi cells in 150 mm plates and treated with either 10 or 320 μM of olaparib for 9 days (> IC90). Fresh media with drug was added every 3 days. Cells that remained attached to the plates were considered persisters. To confirm that the resistant phenotypes were due to adaptation rather than selection of pre-existing resistant clones, we performed two rounds of treatment-recovery of drug-naïve cells (Supplementary Fig 1a). For this, we generated persisters in triplicates with 320 μM (the highest dose used in the escalation experiment) and cells were allowed to recover without drug treatment for 9 days prior to performing viability assays.

### Apoptosis assay

To quantify apoptosis and cell viability of DTPs treated with different doses, we generated persister cells with either 10 or 320 μM of olaparib as previously described. The adherent cells were harvested (0.25% Trypsin/EDTA for 5 min at 37°C), stained with trypan blue and counted manually using hemocytometer or on a Countess II cell counter. Cells were washed twice in cold PBS and stained with DAPI (ThermoFisher) and Annexin V using the Annexin V Apoptosis Detection Kit (BD Biosciences) according to the manufacturer’s instructions. Stained cells were analyzed on a LSR II Flow Cytometer (BD Biosciences). Quantification of apoptotic and viable cells was performed on FlowJo software (FLOWJO).

### CD24 and CD44 cell staining and sorting

The untreated and resistant Kuramochi populations were harvested and stained with DAPI (Cat No. D1306, ThermoFisher), anti-CD24-PE (Cat No. 555428) and anti-CD44-APC (Cat No. 559942) (BD Biosciences). Populations were sorted based on their CD24/CD44 profiles using a Sony SY3200 Highly Automated Parallel cell sorted. Sorted cells were used either for ATAC-Seq or plating for cell viability experiments.

### Generation of patient derived xenografts and drug treatments

All animal experiments were carried out under an approved animal protocol by the Institutional Animal Care and Use Committee (IACUC) of NYU Langone Grossmann School of Medicine. Immunodeficient six-week-old female NOD SCID gamma (NSG) (NOD.Cg-Prkdcscid Il2rgtm1Wjl/SzJ) mice were purchased from Jackson Laboratory and allowed to acclimate to housing conditions. To generate HGSOC (High-Grade Serous Ovarian Cancer) PDX models of PARPi treatments, we used a previously characterized *BRCA2* mutated tumor derived from PDX models (49) (ID: 66799). Approximately 1×10^6^ cells in a 1:1 ratio with Matrigel® Matrix (Corning, USA) were subcutaneously injected with a 27-gauge needle (BD Micro-Fine, USA) into the right lower mammary fat pad on the abdominal dorsal surface. Observation of animals including caliper measurements of tumor volume and body weight continued twice a week. Mice were euthanized with CO2 (confirmed by cervical dislocation) and tumors were excised for further procedures.

*PARPi dose-response:* To study the transcriptional changes that occur upon different dose regimen, the *BRCA2*-mutant HGSOC cells were injected as previously described and mice were randomized into five groups after reaching 100–200 mm^3^ tumor volume. Treatments were performed as follows: a) Vehicle controls (10% dimethylacetamide, 5% Solutol HS15, 85% PBS) (n=2); b) Dose 1 (D1) was 0.13 mg/kg of talazoparib (Selleckchem, S7048) (n=2); c) Dose 2 (D2) was 0.23 mg/kg (n=2); d) Dose 3 (D3) was 0.33 mg/kg (n=2) and e) Dose 4 (D4) was 0.66 mg/kg (5 consecutive days a week, oral gavage). Mice were monitored by routine palpation and tumor sizes measured by caliper. Vehicle tumors were collected at the humane endpoint and talazoparib treated tumors were collected after 50-57 days of treatment (Supplementary Fig. 8a).

*PARPi resistance model:* To establish a drug-resistance model, tumors were treated along three passages. In a first passage, when the PDX tumors reached 100–200 mm^3^ tumor volume, Vehicle (10% dimethylacetamide, 5% Solutol HS15, 85% PBS) or talazoparib (0.0066mg per mouse = 0.33mg/kg (50), oral gavage) was administered for 5 consecutive days a week. Talazoparib treatment caused tumor remission and treatment was interrupted. Tumor sizes were followed until reappearance and treatment was reintroduced. Vehicle and drug treated tumors that did not show further regression upon re-treatment — thus indicating resistance — were collected for scRNA-Seq at the humane endpoint (1500-2000 mm^3^ or up to 20 mm in one dimension). A replicate tumor was implanted in a second passage and maintained under treatment upon tumor detection. The resulting tumor was implanted in a third passage, generating 3 replicates (2 talazoparib treated and 1 vehicle) (Supplementary Fig. 8b).

### Tumor dissociation for scRNA-Seq

Upon arrival to the lab, tumors were minced with a razor blade and enzymatically dissociated in two steps. First, minced samples were incubated with 5 mL of digestion mix 1 (9 mL Advanced DMEM/F12, 1 mL Gentle Collagenase and 100 μL DNase I) at 37°C for 30 min, centrifuged at 1000 RPM for 4 min. After removing the supernatant, samples were incubated with 5 mL of digestion mix 2 (3 mL Advanced DMEM/F12, 2 mL Dispase, 5 mL TrypLE, 100 μL DNase I) at 37°C for 15 min, placed on a 70 μm cell strainer and washed with Advanced DMEM/F12 + 10% FBS. Samples were centrifuged at 1000 RPM for 4 min, incubated with 1 ml of ACK Lysing Buffer (ThermoFisher) for 4 min, washed with PBS + 10% FBS, resuspended in PBS, placed on a 35 μm cell strainer and viability was determined by Trypan Blue staining, which ranged from 65% to 90%. Dissociated cells were collected for scRNA-Seq immediately (resistance experiment) or cryopreserved (dose-response experiment) by resuspending in freezing media (50% FBS, 40% Advanced DMEM/F12 and 10% DMSO) and stored at −80°C. Batches of replicates were processed on the same day. Frozen samples were thawed, washed in media, resuspended in PBS and the cell concentration adjusted for the 10X Genomics single-cell encapsulation protocol following library preparation and sequencing.

### inDrop library preparation and sequencing

Cells from the Kuramochi dose escalation experiment (C, T1-T320) were harvested when they reached confluency at their respective dose treatments and collected for single-cell encapsulation and library preparation using the inDrop platform (69) as previously described (70). Libraries were sequenced on an Illumina NextSeq 500 and reads were processed using a custom inDrop pipeline as described in Baron *et al.* 2020 (70) using the hg38/GRGh38 human genome assembly and the Ensembl 93 transcriptome annotation.

### 10X Genomics library preparation and sequencing

We used the 10X Genomics workflow for scRNA-Seq on cells collected from the experiments depicted in Figure 3 (Kuramochi persisters (P10 and P320) and adapted (T10, T320)), Supplementary Figure 3 (Ovsaho, COV362) and Figure 5 (PDX samples). Cell loading for droplet-based scRNA-Seq and library preparation were performed using the 10X Genomics Chromium platform with the Chromium Single Cell 30 Library & Gel Bead Kit v3.1 following the manufacturer’s instructions. Libraries were diluted to 2 nM and 75bp paired-end sequencing was performed using the Illumina NextSeq 500 and 150-200 million paired reads were generated for each library. Raw reads were processed with the CellRanger v3.1(10X genomics) pipeline (default parameters) with the hg38/GRCh38 genome and Ensembl 93 transcriptome references. For PDX samples, reads were processed with the CellRanger multi-species detection and only cells assigned to ‘human’ were considered for further analysis.

#### Cell hashing for pooled scRNA-Seq

To compare the expression among untreated (C), persister (P10 and P320; two replicates) and the adapted cells (T10 and T320), we used the Cell Hashing method (71) to tag cells from different conditions based on the expression of surface markers (TotalSeqA, Hashtag oligos (HTOs) 1-5, Biolegend). Five samples from each condition were pooled into a single cell suspension in equal amounts and loaded on the 10X Genomics Chromium Controller with Single Cell 3’ v3.1 system and libraries prepared according to the manufacturer’s protocol. For the hashtag oligos, 1ul of HTO PCR additive primer was added to the 10X cDNA amplification step, and the supernatant from the 0.6X cDNA cleanup was kept and processed according to the Cell Hashing protocol, with 14 PCR cycles and a 1.2X cleanup after the PCR. The same procedure was applied to pool cells from Ovsaho (C, T5-T40; HTOs 1-5) and COV362 (C, T5-T40; HTOs 6-10) experiments. Sample demultiplexing was performed using the CITE-seq-Count pipeline (v1.4.3, https://github.com/Hoohm/CITE-seq-Count) and Seurat (16).

### ATAC-Seq sample preparation and sequencing

ATAC-Seq samples were prepared in duplicates using a modified version of the OMNI-ATAC protocol (72). Briefly, cells grown in tissue culture were trypsinized, stained for expression of CD44 and CD24 and sorted as previously described. Briefly, approximately 50,000 cells per condition were used for nuclei extraction using the ATAC-Seq resuspension buffer as previously described (72). Nuclei were resuspended in 50 μl of transposition mix (25 μl 2× TD buffer, 2.5 μl transposase (100 nM final), 16.5 μl PBS, 0.5 μl 1% digitonin, 0.5 μl 10% Tween-20, and 5 μl water). Transposition reactions were incubated at 37°C for 40 min in a thermomixer with shaking at 1,000 r.p.m. Reactions were cleaned up with Zymo DNA Clean and Concentrator 5 columns. ATAC-Seq libraries were amplified using Ad2_noMx (universal forward) and Ad2.x (indexed reverse). Libraries were diluted to 2nM and 75bp paired end sequencing was performed using the Illumina NextSeq 500/550. Between 50-200 million paired reads were generated for each library.

### Single-cell data analysis

*Filtering:* Raw gene expression matrices were TPM normalized (scale factor of 10^4^) and log_2_ transformed. For each cell, the number of transcripts, genes expressed, and the proportion of transcripts derived from mitochondrial and ribosomal genes were determined. For each dataset, cells with transcript number lower than 500 or higher than 2 standard deviations, with less than 500 genes detected and with higher than 20% of transcripts corresponding to mitochondrial genes were excluded from further analysis. Genes expressed in less than 1% of the cells in each dataset were removed.

#### Cell clustering and differential gene expression

For all samples we observed that differences due to cell cycle gene expression was a major source of variation. To mitigate this effect for the interpretation of cell states, we used the Seurat function *CellCycleScoring* to assign cells in G1, S and G2/M phases. We then considered the G1 cells for further analyses. Dimensionality reduction, clustering and differential expression were performed using the Seurat R package. To control for unwanted sources of variation, the number of UMIs and the fraction of mitochondrial and ribosomal UMIs were regressed out during data scaling. For the individual analysis of the Kuramochi datasets (C-T320) we used the DrImpute method (k parameter = 5:10) (73) before clustering. Highly variable genes were identified using the Seurat *FindVariableGenes* function (default parameters) and these were used to compute the Principal Components (PCA) following Uniform Manifold Approximation and Projection (UMAP) using the top 5 to 10 PCs (cutoffs determined by the elbow method). Louvain clustering was performed using the *FindClusters* function using the selected PCs with resolution set to 0.3 to 0.6. For the analysis of PDX dose-response samples, we observed a batch effect driven mostly by one of the vehicle samples and we used the *fastMNN* method (74) for correction of the aggregated datasets before clustering. Differential gene expression across clusters was performed using the *FindMarkers* function using the MAST test (logfc.threshold=0.25, min.pct=0.2, min.cells.genes=10).

#### Defining cell states across resistant populations

To study shared cellular states across the Kuramochi resistant populations, we first assessed the differentially expressed genes among the cell clusters (i.e., cell subpopulations) within each sample separately. The top 100 most differentially expressed genes (ranked by their adjusted p-values) in each cluster were selected to generate a non-redundant list of genes (n = 1511). These genes were used to build an average expression matrix of each cell cluster across all samples. A correlation matrix (Spearman’s correlation coefficients) across these average profiles was used for average linkage hierarchical clustering and the cell states were defined based on the cluster structure (k = 5).

#### Gene module identification

To identify gene expression programs across the Kuramochi lines from C to T320, we implemented the Neftel *et al.* 2019 approach (17), with some modifications. First, we aggregated all G1 cells (non-imputed dataset) from all conditions and performed an average linkage hierarchical clustering using the 1000 most highly variable genes (defined by the *FindVariableGenes* function) on the centered dataset using one minus the Pearson correlation as the distance metric, as proposed previously (17). To evaluate all possible clusters of cells that have a potential gene expression signature, we recovered all clusters excluding by size (less than 30 cells and no more than 85% of all cells) and by signal of differential expression. Each cluster of cells was evaluated for a signature of differential expression relative to the remaining cells using the *FindMarkers* function as described above. Clusters with a signature of differentially expressed genes were retained according to the following criteria: genes expressed > 1.8 average fold change, adjusted p-value < 0.001 and with more than 50 genes matching these criteria. Gene signatures (i.e., gene sets) were then hierarchically clustered based on their Jaccard index and, for pairs with similarity above 0.75, the signature with the largest number of genes was considered. To integrate the gene signatures into gene modules, clusters with less than 3 signatures were excluded. The genes from all signatures that constitute a particular cluster were merged and ranked based on the ratio of their average expression in cells from a particular cluster relative to their average expression in cells from the other clusters. To assess the significance of gene overlap across modules and differentially expressed gene lists from other analyses, p-values were computed with Fisher’s Exact tests implemented in the *GeneOverlap* R package. Cells were scored by their module expression (top 100 genes excluding mitochondrial and ribosomal genes) using Seurat’s *AddModuleScore* method and z-scored for visualization. To assess the significance of overlap among genes in modules and genes in CNA (Copy Number Alteration) regions, permutation tests (10, 000) were performed using the *regioneR* package (75).

#### Identification of genes with dose-response patterns

Using the PDX datasets from the dose-response experiments, we identified genes with dose-response patterns by first performing pairwise differential expression across all treated conditions versus vehicles for each batch of replicates independently. Genes differentially expressed in both of the replicates were considered. A non-redundant list of these differentially expressed genes was used to generate an average expression matrix across conditions. The samples were ordered from low to high doses (V, D1, D2, D3, D4) and the genes were clustered using the *Mfuzz* package (76) setting the number of clusters to 7 and m parameter = 1.8. Genes with membership score higher than 0.3 were included and those with similar profiles of decreasing or increasing expression relative to high (or low) doses in both replicates were aggregated.

### Single-cell inferred copy number alterations (CNAs)

Large-scale chromosomal alterations from scRNA-Seq datasets were inferred by the *inferCNV* v1.4.0 package. Untreated samples were used as a reference for CNA inference in the treated samples. The default options were used except for a cutoff of 0.1 for the minimum average read counts per gene and clustering by groups was set based on the cluster annotations determined from the transcriptome analyses. The output consisting of a denoised modified expression matrix as genes ordered by chromosomal location was used for heatmap visualizations with *ComplexHeatmap* (*77*) in R.

### Gene set enrichment analysis

Gene set enrichment using the hallmark annotations (MSigDB) (78), KEGG (79) and Reactome (80) pathways was performed using the R package *fgsea*. Differential expression between groups was determined using the MAST algorithm implemented in Seurat and the genes were ranked using the sign of the average log fold change multiplied by the −log10 of p-value. Ranked lists were used as inputs for the *fgsea* package with a minimum gene set size of 10 and a maximum of 500, and significance assessed by 10,000 permutations. Terms were considered significantly enriched at p-values < 0.05.

### ATAC-Seq data processing and analysis

#### Peak detection

Adapter sequences were removed using Trim Galore (https://github.com/FelixKrueger/TrimGalore) (-q 20, - --length 30). Filtered reads were aligned to the human genome (hg38/GRCh38) with Bowtie2 (81) (--very-sensitive, -k 10, -X 2000). After removing reads mapping to mitochondrial DNA and duplicate reads removed with Picard Tools MarkDuplicates (http://broadinstitute.github.io/picard/), only properly mapped pairs assigned by SAMTools (82) were included for further analysis. To avoid peak detection bias due to sequencing depth differences, read depth across samples was downsampled with SAMTools to match the lowest depth. Peak calling was performed with MACS2 (83) and blacklisted regions were excluded (84). Reproducible peaks between replicates for each sample were determined using the IDR framework (https://github.com/nboley/idr) and peaks with high reproducibility (IDR score < 0.05) were considered. A consensus peak set across all samples was constructed by merging the reproducible peaks from each sample using HOMER v4.1 (85). Finally, a read count table for the consensus peaks was generated using deepTools 3.1 (86).

#### Differential peak accessibility and motif analysis

The count table of consensus peaks was normalized and differentially accessible peaks were calculated using *DESeq2* (87). Differentially accessible peaks relative to the untreated sample (C) were considered as adjusted p-value < 0.001 and fold change ≥ 3. The top 2000 most variable peaks were used for Principal Component Analysis (PCA). To visualize the peak signal across samples, ‘Zavit’ (48,88) was used to sort the mean peak signal (between replicates) across samples. Motif enrichment analysis on the differentially accessible peaks was performed using HOMER’s ‘findMotifsGenome’ for known transcription factor motifs. To assess the significance of the overlap between the genomic regions of the differentially accessible peaks and the regions that encompass copy-number alterations (CNAs determined by WES), permutation tests (1,000 permutations) were performed using the *regioneR* package. To verify the influence of CNAs on the motif enrichment patterns, the motif analysis was repeated by removing the more or less differentially accessible peaks that overlap with gained or deleted copy-number regions, respectively.

### Whole exome sequencing (WES) and analysis

Genomic DNA extraction for whole exome sequencing was performed using Qiagen Blood and Cell DNA kit (Catalog no: 13323). The extracted DNA from representative samples (C, T10, T40 and T320) was sent to the Broad Institute facility and processed according to the in-house pipeline. Briefly, 125ng in 50μL was prepared for sequencing using the KAPA Hyper Prep Kit. Hybridization and capture were performed using the relevant components of IDT’s XGen hybridization and wash kit and following the manufacturer’s suggested protocol. Sequencing was performed using Illumina Novaseq.

Sequencing results were demultiplexed and converted to FASTQ format using Illumina bcl2fastq software. The FASTQ files were processed using the Seq-N-Slide pipeline (https://doi.org/10.5281/zenodo.5550459). Adapter sequences were trimmed with Trimmomatic (89) and then aligned to the human reference genome (build hg38/GRCh38) using the Burrows-Wheeler Aligner with the BWA-MEM algorithm (90). Low confidence mappings (mapping quality < 10) and duplicate reads were removed using Sambamba (91). Further local indel realignment and base-quality score recalibration were performed using the Genome Analysis Toolkit (GATK) (92). Copy number profiles were calculated using Control-FREEC (93) with untreated samples as the matched controls. ANNOVAR (94) was used to annotate variants with genomic context such as functional consequence on genes and identify presence in public variant databases gnomAD and COSMIC.

## Acknowledgements

We thank Rich White, Markus Schober, Maria D. Vibranovski, Andrei Rozanski and the members of the Yanai lab for helpful discussions. We thank the lab of Benjamin Neel for providing the cell lines used in this study. This work was supported by the following NIH grants: P50 CA225450 (to I.Y.), R01 LM013522 (to I.Y.), R21 CA264361 (to I.Y.), GM126573 and F30 CA257400 (to D.B.).

## Author contributions

GSF conceived and performed all of the *in vitro* experiments and analysis. GSF and IY designed most of the experiments and wrote the manuscript. MB performed the scRNA-Seq for the dose-escalation experiment. BK, TL and DAL designed the *in vivo* resistance experiment. BK and SM performed the *in vivo* resistance experiment. SM performed all of the mouse work. GSF, BK, SM, TL,and IY designed the *in vivo* persistence experiment. GSF, MP, BK and SM performed the *in vivo* persistence experiment with assistance from DB. MP assisted with the *in vitro* persister experiments, and led the single-cell processing of the *in vivo* persistence experiment. GSF, AR and IY designed the ATAC-Seq experiments. AR performed the ATAC-Seq experiments, 10X library preparations and DNA extraction. GA and FK assisted with scRNA-Seq for Kuramochi persister, Cov362 and Ovsaho. KHT and AP assisted with additional *in vitro* experiments. MB, MP, AR, BK and DB assisted with result interpretations. ID contributed with whole-exome analysis. IY oversaw the coordination of the project. All authors commented on the manuscript.

